# omicsGMF: a multi-tool for dimensionality reduction, batch correction and imputation applied to bulk- and single cell proteomics data

**DOI:** 10.1101/2025.03.24.644996

**Authors:** Alexandre Segers, Cristian Castiglione, Christophe Vanderaa, Lennart Martens, Davide Risso, Lieven Clement

**Affiliations:** Department of Applied Mathematics, Computer Science and Statistics, Ghent University. Ghent, Belgium; Center for Medical Genetics Ghent, Ghent University and Ghent University Hospital. Ghent, Belgium; Bocconi Institute for Data Science and Analytics, Bocconi University. Milan, Italy; Department of Biomolecular Medicine, Ghent University. Ghent, Belgium; VIB-UGent Center for Medical Biotechnology, VIB. Ghent, Belgium; Department of Statistical Sciences, University of Padova. Padova, Italy

## Abstract

The unprecedented speed and sensitivity of mass spectrometry (MS) unlocked large-scale applications of proteomics and even enabled proteome profiling of single cells. However, this fast-evolving field is hindered by a lack of scalable dimensionality reduction tools that can compensate for substantial batch effects and missingness across MS runs. Therefore, we present omicsGMF, a fast, scalable, and interpretable matrix factorization method, tailored for bulk and single-cell proteomics data. Unlike current workflows that sequentially apply imputation, batch correction, and principal component analysis, omicsGMF integrates these steps into a unified framework, dramatically enhancing data processing and dimensionality reduction. Additionally, omicsGMF provides robust imputation of missing values, outperforming bespoke state-of-the-art imputation tools. We further demonstrate how this integrated approach increases statistical power to detect differentially abundant proteins in the downstream data analysis. Hence, omicsGMF is a highly scalable approach to dimensionality reduction in proteomics, that dramatically improves many important steps in proteomics data analysis.

---

Recent technical advancements in mass spectrometry (MS) have enabled the proteomewide characterization of biological samples with unprecedented speed and sensitivity. These developments have facilitated the application of proteomics in large-scale clinical studies (e.g., [1, 2]) while simultaneously enabling the characterization of thousands of proteins at the single-cell level [3, 4]. However, this increased throughput is accompanied by substantial technical batch effects and a high prevalence of missing data, primarily due to the extensive number of MS runs required for such applications [1–5]. Together, these challenges hinder data exploration, normalization, and subsequent differential analysis.

Typically, a fundamental initial step in such large-scale proteomics (LSP) data analysis involves dimensionality reduction (DR), which facilitates data visualization and the extraction of meaningful insights from their data. Furthermore, DR is crucial in single-cell proteomics (SCP) for downstream analyses, including cell clustering, denoising, and trajectory inference [6]. However, DR in SCP presents significant challenges due to the high degree of missing data, which ranges from 50% to 90% [7]. Standard principal component analysis (PCA) methods are not applicable in the presence of missing values. To address this limitation, extensions of conventional PCA, such as NIPALS [8, 9] and expectation-maximization PCA [10], have been proposed. Nevertheless, these approaches suffer from numerical instability and high computational complexity, making them unsuitable for large-scale SCP datasets [7]. Moreover, they are unable to effectively correct for known batch effects, further restricting their practical applicability in proteomics.

Consequently, conventional workflows continue to rely on imputation of missing values prior to DR. However, not all missing values arise from the same underlying mechanism. Specifically, missingness may occur due to variations in ionization efficiency, ion competition, or computational limitations (e.g., unreliable identification), which are independent of peptide abundance or the abundance of other peptides. These types of missingness are all classified as missing completely at random (MCAR). In contrast, certain peptides may be absent because they are not present in the cell or sample, or because their abundance falls below the detection limit, a phenomenon categorized as missing not at random (MNAR). Given these differences, distinct imputation strategies are required for MCAR and MNAR peptide-spectrum matches (PSMs) [11]. Indeed, the application of imputation methods designed to address only one type of missingness can significantly alter the distribution of protein-level intensities [12].

Moreover, the prevalence of MCAR and MNAR is largely influenced by the data acquisition strategy, as well as by the specific characteristics of peptides and proteins, making the imputation process particularly challenging. To address this issue, more sophisticated methods have been developed, leveraging machine learning approaches [13] or explicitly estimating the proportions of MCAR and MNAR values to inform the selection of an appropriate imputation strategy [14]. However, these methods are primarily designed for label-free bulk proteomics with data-dependent acquisition strategies, whereas labeled approaches and data-independent acquisition techniques are much more commonly employed in LSP and SCP to enhance proteome coverage. In Fig. 1 and Supplementary Fig. 1, we illustrate on three example datasets that employing PCA for visualization after conventional imputation approaches is mainly driven by the large number of missing values rather than by biological sources of variability. Panel B highlights another limitation: PCA does not remove batch effects, which are known to be major sources of variability in LSP and SCP experiments [5, 15]. Consequently, an additional preprocessing step is required to eliminate batch effects, resulting in lengthy and complex analytical workflows that necessitate proficiency with multiple computational tools.

**Fig. 1.**
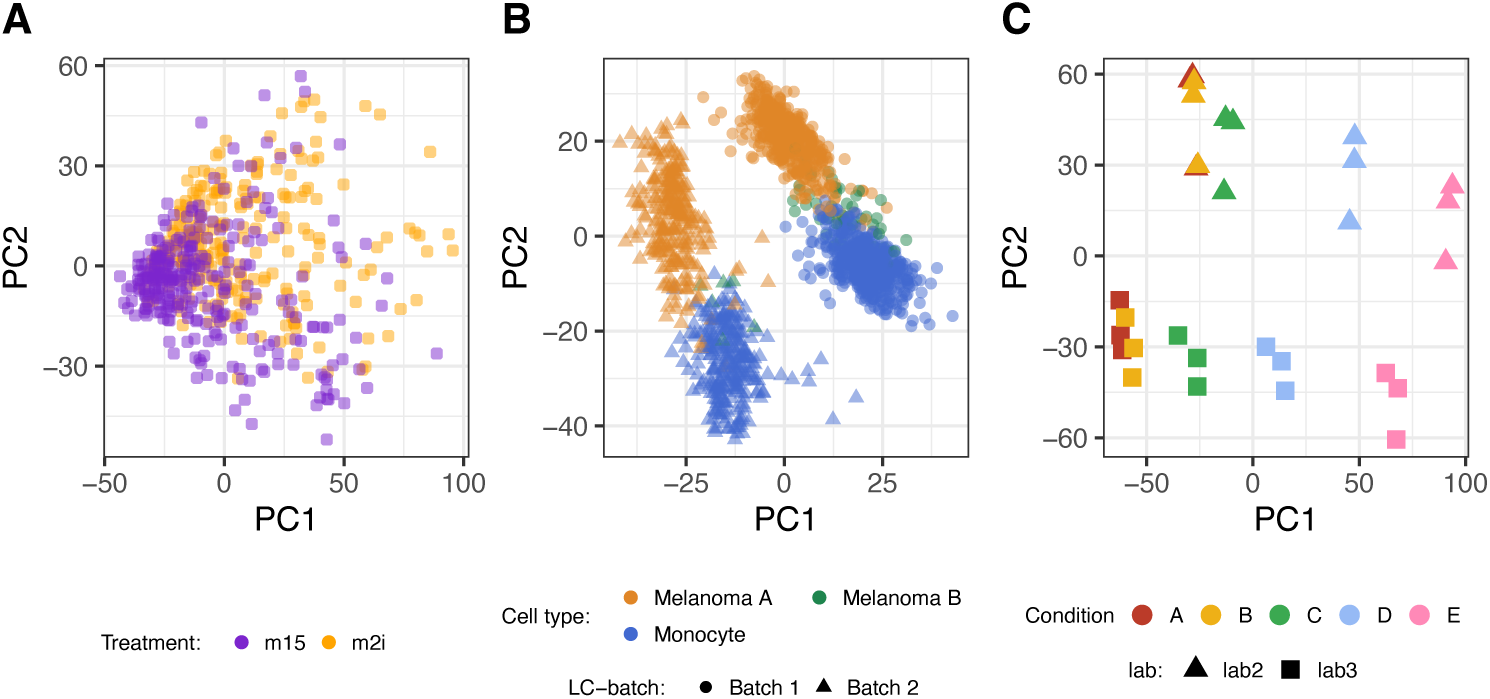
Issues with conventional multistep workflows for dimensionality reduction upon imputation. Panel A shows data from the label-free, single cell Petrosius study [4] where the clusters of mouse-embryonic stem cells treated with and without inhibitor largely overlap, while the treatment is expected to change the proteome considerably. In Panel B a PCA-plot is made for the labeled single cell Leduc dataset [3] highlighting that batch effects are the main source of variability. It overwhelms the variability associated with the melanoma B subpopulation and renders the first dimensions obsolete for clustering cell-types. Panel C shows data from the label-free, bulk CPTAC spike-in study [16] with 48 human UPS proteins that were spiked in at five different concentrations in a yeast background. The experimental conditions with the lowest spike-in concentrations (Condition A and B) cannot be separated by the conventional multi-step workflow. All low dimensional visualizations were obtained with state-of-the-art CF-imputation [13] followed by PCA.

More critically, the order in which batch correction and imputation are performed also has a substantial impact on downstream analyses [17], underscoring the necessity for approaches that perform batch correction and imputation, simultaneously. In this context, scPROTEIN [18] represents an initial attempt to address both batch effects and missing data. However, its batch correction is limited to a single factor, lacking the flexibility to account for multiple technical artifacts and other confounding variables. Given the importance of adjusting for these sources of variation in LSP and SCP experiments, more comprehensive solutions are needed to enhance the robustness and accuracy of proteomic data analysis.

Therefore, the development of novel dimensionality reduction methods that can simultaneously account for batch effects, handle missing values, and scale efficiently to the increasing data volumes in LSP and SCP experiments is essential to push the field forward. In this work, we develop a user-friendly package, omicsGMF, that leverages the power of our novel sgdGMF framework, stochastic gradient descent for generalized matrix factorization [19], with the omics data infrastructure of the Bioconductor ecosystem to develop streamlined workflows that specifically addresses the unique challenges posed by large-scale (single-cell) proteomics datasets. omicsGMF is designed to correct for known sample- or feature-level covariates, accommodate missing values, and optimize its parameters through an efficient adaptive stochastic gradient descent algorithm leveraging minibatch subsampling, partial parameter updates, and exponential gradient averaging. These computational strategies provide substantial efficiency gains over traditional matrix factorization approaches that support missing data. As a result, omicsGMF effectively addresses the three primary challenges of dimensionality reduction in the LSP and SCP contexts, while preserving the interpretability of conventional PCA by maximizing a Gaussian likelihood function.

We first demonstrate that omicsGMF provides superior dimensionality reduction for both TMT-labeled and label-free proteomics data by benchmarking it against commonly used workflows that involve separate imputation, batch correction, and principal component analysis (PCA), as well as against the single-step method scPROTEIN [18]. Next, we illustrate how the final parameter estimates from omicsGMF can be leveraged for imputation of missing values and evaluate its performance relative to traditional imputation methods, including k-nearest neighbors (KNN) imputation [20], quantile regression imputation of left-censored data (QRILC) [21], zero or minimum imputation, and more recent deep-learning-based approaches [13]. Finally, we present a case study with known ground truth, demonstrating that imputation performed by omicsGMF yields superior results for downstream differential abundance analysis.

## Results

omicsGMF leverages our novel sgdGMF framework [19] to develop innovative workflows for information extraction that tackle key challenges in proteomics data analysis. Indeed, omicsGMF can concentrate the leading sources of variability in a limited number of dimensions, while accounting for missing values and known covariates such as treatment and batch effects (Fig. 2). Consider *n* samples (bulk samples or single cells) and *J* features (PSMs, peptides or proteins). The matrix with normalized intensities (**Y**) with entries *y_ij_* of sample *i* (*i* = 1*, …, n*) and feature *j* (*j* = 1*, …, J*) can then be modeled in terms of known sample- and feature-level covariate matrices, **X** and **Z**, such as cell types, experimental batches and quality control measures. Further, consider the unknown sample-level latent covariate matrix **U**, which accounts for unknown variation as in RUV [22] and ZINB-WaVe [23]. These primal directions of unknown (biological) variation represent the samples in a reduced dimensionality and are often used for downstream analyses such as visualization or clustering of samples. The parameters of the known sample and feature covariates, and the unknown latent covariates are ***B***, **Γ** and **V**, respectively.

**Fig. 2.**
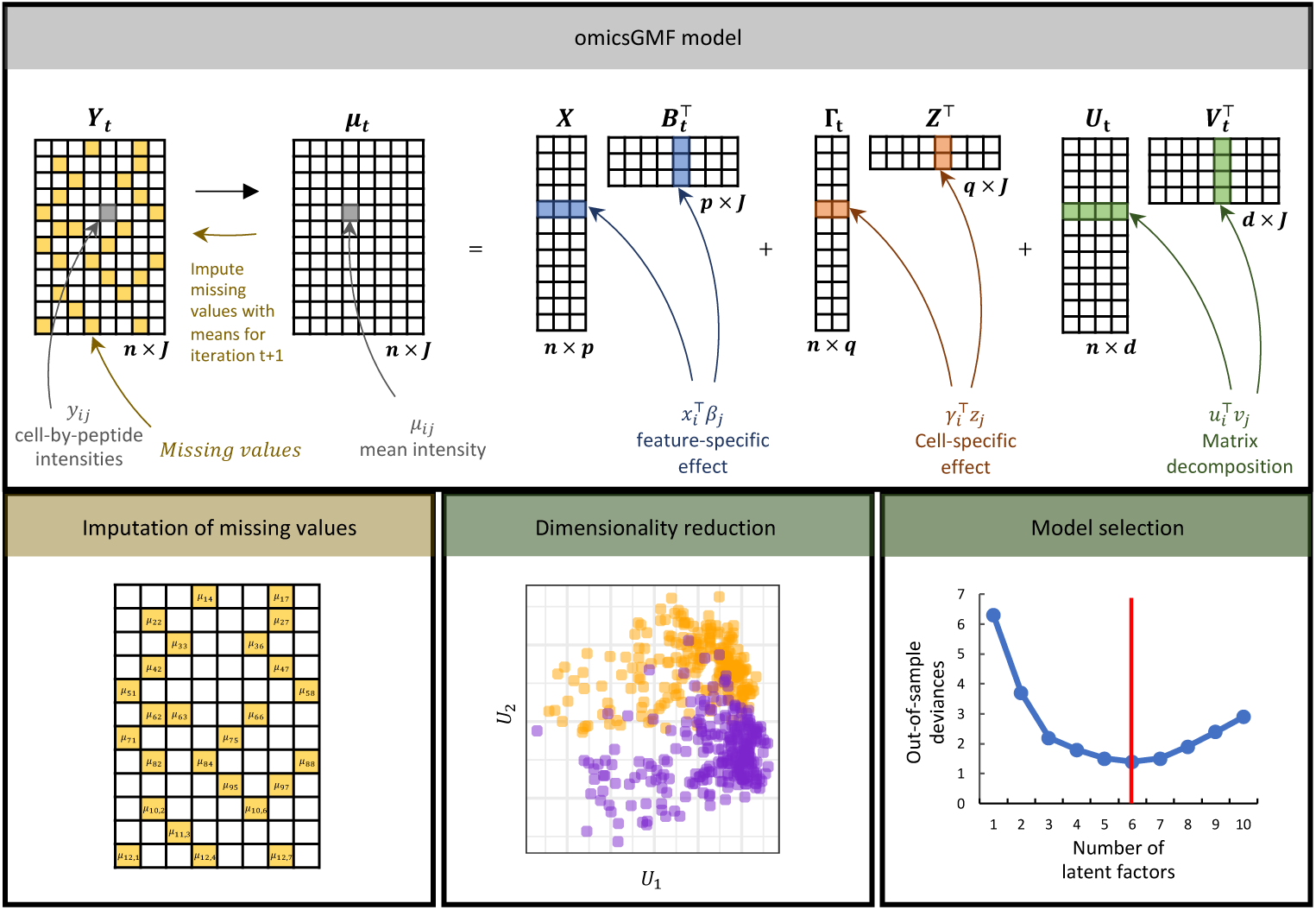
Schematic overview of the omicsGMF model for the Gaussian Model family. Y is modeled in function of known sample-level covariates X, feature-level covariates Z, latent factors U and their loadings V. omicsGMF iteratively estimates the parameters ***B***, L, U and V. omicsGMF addresses missing values by re-imputing them in each iteration with their current mean ***µ****_t_*. The latent factors have a similar interpretation as principal components upon correcting for known covariates, and thus allow for dimensionality reduction and visualization. omicsGMF can also provide the imputed values upon convergence, which are useful for downstream applications. Furthermore, omicsGMF allows for model selection that can guide the user for choosing the number of latent factors and known covariates to be included in the model. More details can be found in the Methods section.

omicsGMF estimates all the unobserved parameters **U**, ***B***, ***L*** and **V**. Since both **U** and **V** are unknown, a closed-form solution is not available. Therefore, omicsGMF uses iterative regression of **Y** on [**X**, ***L***^^^, **U**^^^ ] followed by regression of **Y** on [***B***^^^ , **Z**, **V**^^^ ] to estimate all the parameters. Here, the hat notation indicates the current estimate of the corresponding parameters. Due to the extensive number of missing values, these linear regressions are not possible without imputing such unobserved data. Therefore, omicsGMF iteratively imputes the missing values with the estimated mean of the prior iteration. The advantages are threefold. First, the imputed values account for known sample- and feature-level covariates, as well as the latent structure, resulting in accurate imputed intensities even when dealing with significant technical effects. Second, it avoids complicated multi-step workflows and performs imputation and correction for technical effects simultaneously. Therefore, the user does not suffer from different results that rely on the order in which batch correction and imputation are conducted [17, 24]. Third, it is an all-in-one workflow that maximizes a Gaussian likelihood, rendering its parameter estimates, latent variables, and imputed values easily interpretable. Additionally, the latent covariates **U** used for dimensionality reduction and visualization have a similar interpretation to conventional PCA.

A critical issue in conventional algorithms and pipelines is their lack of a principled way to estimate the optimal number of latent factors, *d*, in the model. Remarkably, the omicsGMF framework implements several model selection methods relying on scree plot investigation, eigenvalue thresholding, minimization of information-based criteria, e.g., AIC and BIC, and minimization of the cross-validation error to evaluate the out-of-sample error on subsets of holdout values masked and estimated as if they were missing observations. This results in comprehensive and easy-to-use model selection for the user, allowing for optimal downstream analyses. Moreover, omicsGMF uses minibatch subsampling, partial parameter updates and exponential gradient averaging to obtain computational advantages over existing methods that can deal with missing values. We refer to the Methods section for an overview of the omicsGMF estimation and imputation processes, and an extensive overview of the approach’s minibatch sub-sampling, partial parameter updates and exponential gradient averaging is provided here [19].

The benchmarking of omicsGMF versus other state-of-the-art tools consists of three parts. We first show that omicsGMF can accurately visualize proteomics data while correcting for known confounders such as batch effects. We then compare omicsGMF imputation to other state-of-the-art imputation tools in a benchmark that includes missing values that are missing completely at random (MCAR) as well as missing values due to low abundance, i.e., missing not at random (MNAR). We also evaluate the distributions of imputed intensities in a dataset with known spike-in concentrations of human proteins. Finally, we compare the performance of different imputation strategies in downstream differential abundance analyses. These benchmarks show that omicsGMF is an all-in-one tool that outperforms current complex multistep workflows for dimensionality reduction and visualization of proteomics data, with superior performance in downstream analyses while remaining fast, scalable and easy to use.

### Dimensionality reduction and visualization of proteomics data

We demonstrate that the latent factors estimated by omicsGMF can be used for the visualization of high throughput data with a similar interpretation as PCA upon correction of known covariates. We compare the visualizations provided by omicsGMF to conventional workflows that first impute missing data, and subsequently use ordinary PCA on the imputed data matrix. This is done for the label-free single-cell Petrosius dataset [4] consisting of 525 cells and 4435 peptides with 58.5% missing values, the TMT-labeled single-cell Leduc dataset [3] consisting of 1508 cells and 6280 peptides with 61.8% missing values, and the bulk label-free CPTAC study [16] where 48 human UPS proteins were spiked in at five different concentrations in a yeast proteome background. In the latter experiment, three samples for each concentration were analyzed in three different labs, resulting in a total of 45 MS-runs. In the main text we only show the results for a multi-step workflow with a PIMMS neural network-based collaborative filtering (CF) [13] and K-Nearest-Neighbors (KNN) imputation followed by PCA. We refer the reader to supplementary information for results on PIMMS denoising autoencoder (DAE) [13], PIMMS variational autoencoder (VAE) [13], QRILC [21], and zero or minimum imputation imputation. The Supplementary Figures also include data exploration with NIPALS [9], which performs PCA by ignoring missing values, and therefore does not require a prior imputation step.

Visualization upon dimensionality reduction of the Petrosius dataset (Fig. 3, Panel A) shows that omicsGMF provides a better separation between treated and untreated mouse stem cells. CF imputation prior to PCA does not show a good visual separation according to treatment in the space defined by the first two PC’s. The same holds for the other bespoke proteomics imputation methods DAE, VAE, QRILC, and zero and minimum imputation (Supplementary Fig. 2). In fact, these are also out-competed by basic KNN-imputation. Interestingly, the first two principal components of omicsGMF have much lower correlation with missing values, suggesting that omicsGMF suffers less from missing data artifacts (Supplementary Fig. 3 and 4).

**Fig. 3.**
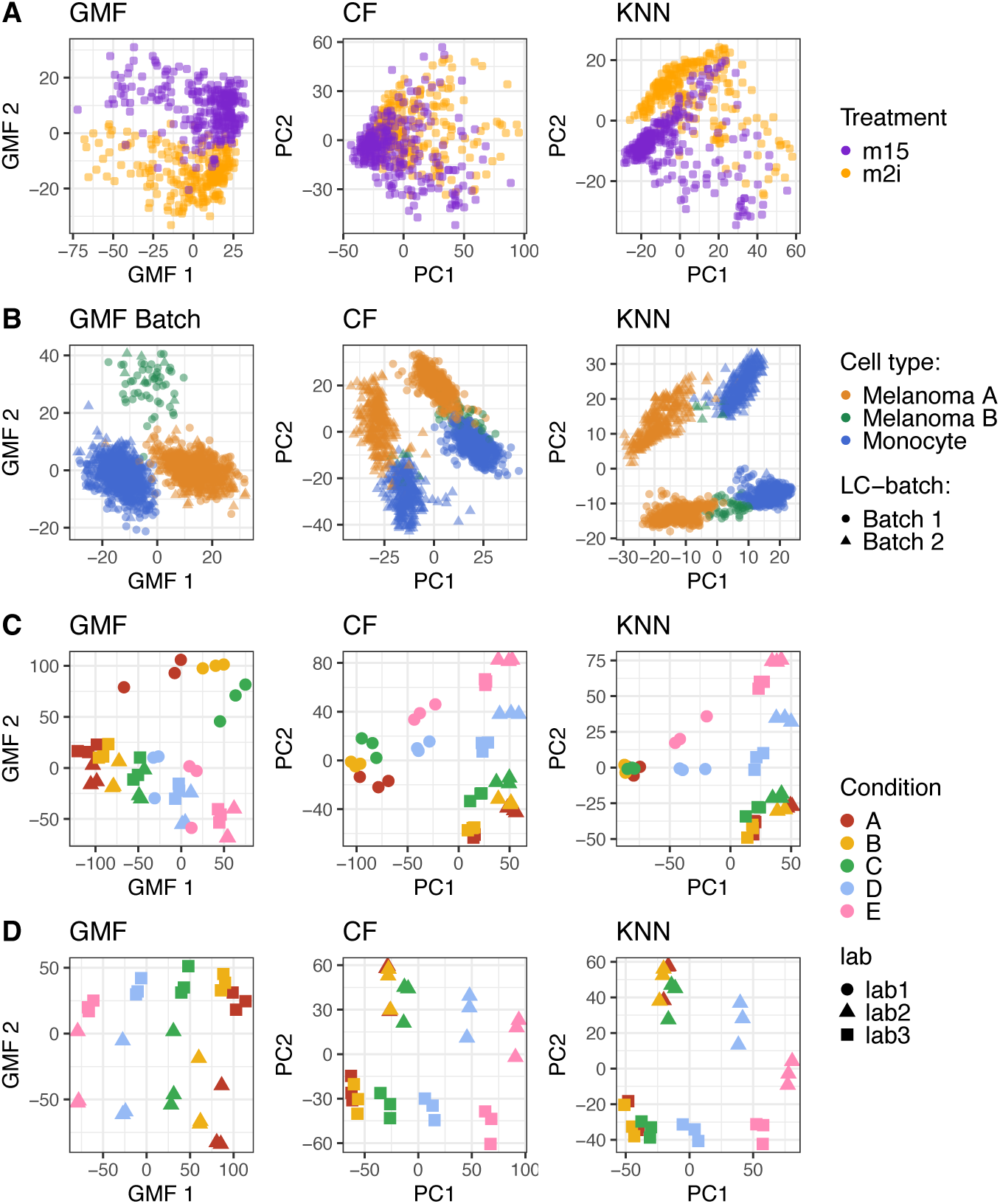
Low-dimensional visualization of proteomics data. omicsGMF estimates latent factors that have a similar interpretation as regular PCA. These can be used for a low dimensional visualization of proteomics data and are compared to PCA plots after CF and KNN imputation of missing data. Panel A shows the results for the Petrosius [4] dataset, colored by inhibitor treatment. Panel B shows different cell-types from the Leduc [3] dataset. Here, omicsGMF directly accounts for known batch effects, resulting in a better representation of the biological signal compared to PCA after CF and KNN-imputation. Panel C and D show CPTAC data [16] from all labs, and upon exclusion of Lab 1, respectively (Lab 1 was known to suffer from ionization issues). Samples are colored by the spike-in concentration of human proteins, with A the lowest spike-in concentration, and E the highest spike-in concentration. Distinct marker shapes indicate the different labs.

omicsGMF can also directly account for known covariates, such as batch effects, which we illustrate for the TMT-labeled Leduc dataset (Fig. 3, Panel B; Supplementary Fig. 5). Indeed, Leduc et al. multiplexed up to 14 cells in each TMT run. Upon correcting for the run effect omicsGMF uncovers additional biological variability corresponding to a previously reported subpopulation of melanoma cells [3]. Imputation of missing values followed by PCA cannot directly control for the technical run-to-run variability, resulting in a dimensionality reduction where the first two latent variables are mainly driven by this strong batch effect. Hence, conventional PCA-based workflows with imputation methods require an additional step for batch removal prior or post imputation. NIPALS, however, which conceptually does not require imputation of missing values, also cannot account for known batch effects, and therefore suffers from the same caveats. When using a more complicated multi-step workflow involving imputation, batch correction, and PCA, similar visualizations as with omicsGMF can be obtained for CF, DAE, VAE, NIPALS and KNN, while QRILC, zero and minimum imputation clearly display imputation artifacts in their low dimensional visualizations (Supplementary Fig. 6).

**Fig. 4.**
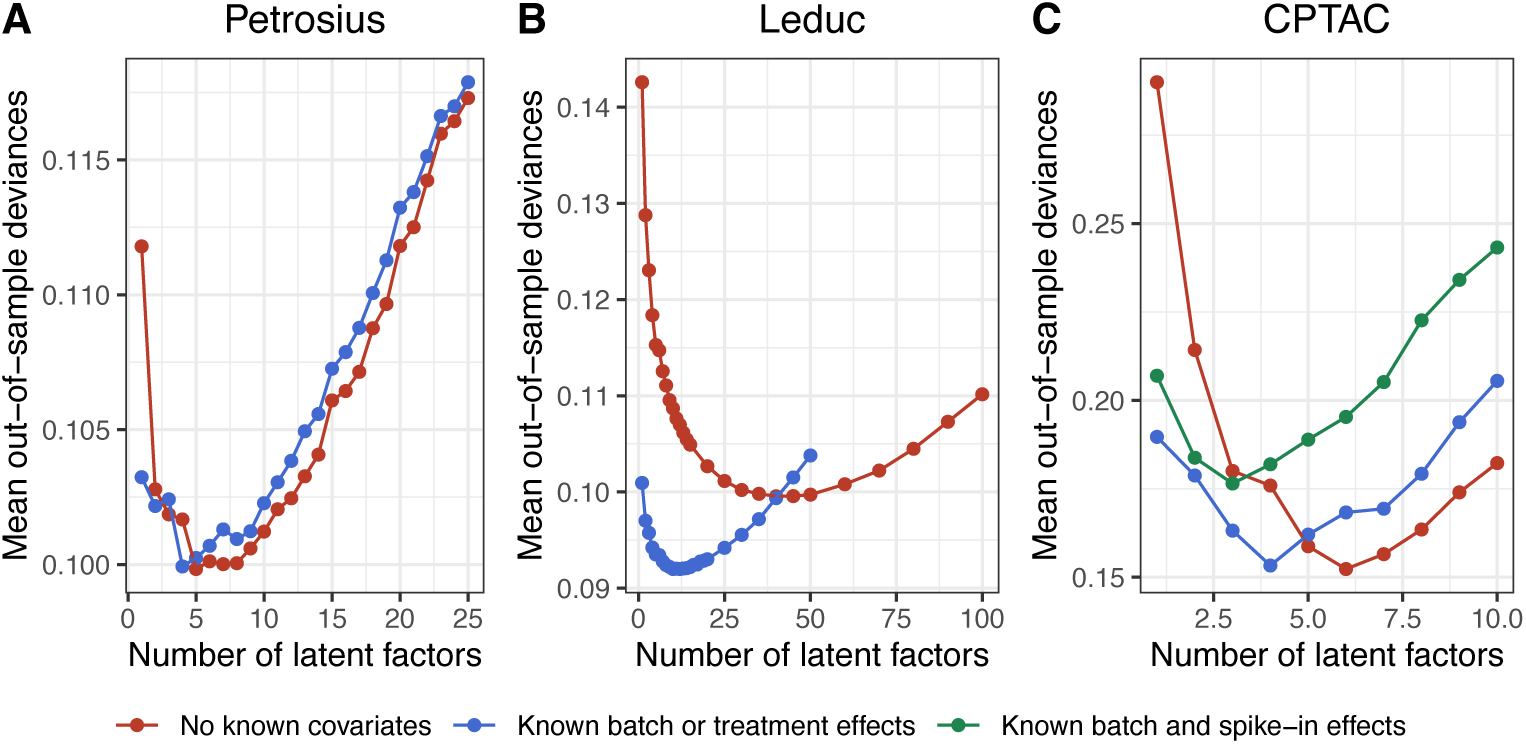
Cross-validation with omicsGMF allows for comprehensive selection of the number of latent factors for dimensionality reduction. Each panel shows the mean of the out-of-sample deviances over three cross-validation folds in function of the number of latent factors included in the model. In each fold, 30% of the values are masked for out-of-sample prediction. Panel A shows the cross-validation results for the Petrosius dataset [4] with and without accounting for the treatment effect (one dummy variable). Panel B shows the cross-validation results for the Leduc dataset [3], with and without correcting for the known batch-effect associated to multiplexing cells in the same run (142 dummy variables). Panel C shows the cross-validation results for the CPTAC data [16], considering all three labs. Results are shown for omicsGMF without known covariates, accounting for the lab effects (two dummy variables), and accounting for both the lab (two dummy variables) and spike-in concentration effects (four dummy variables).

**Fig. 5.**
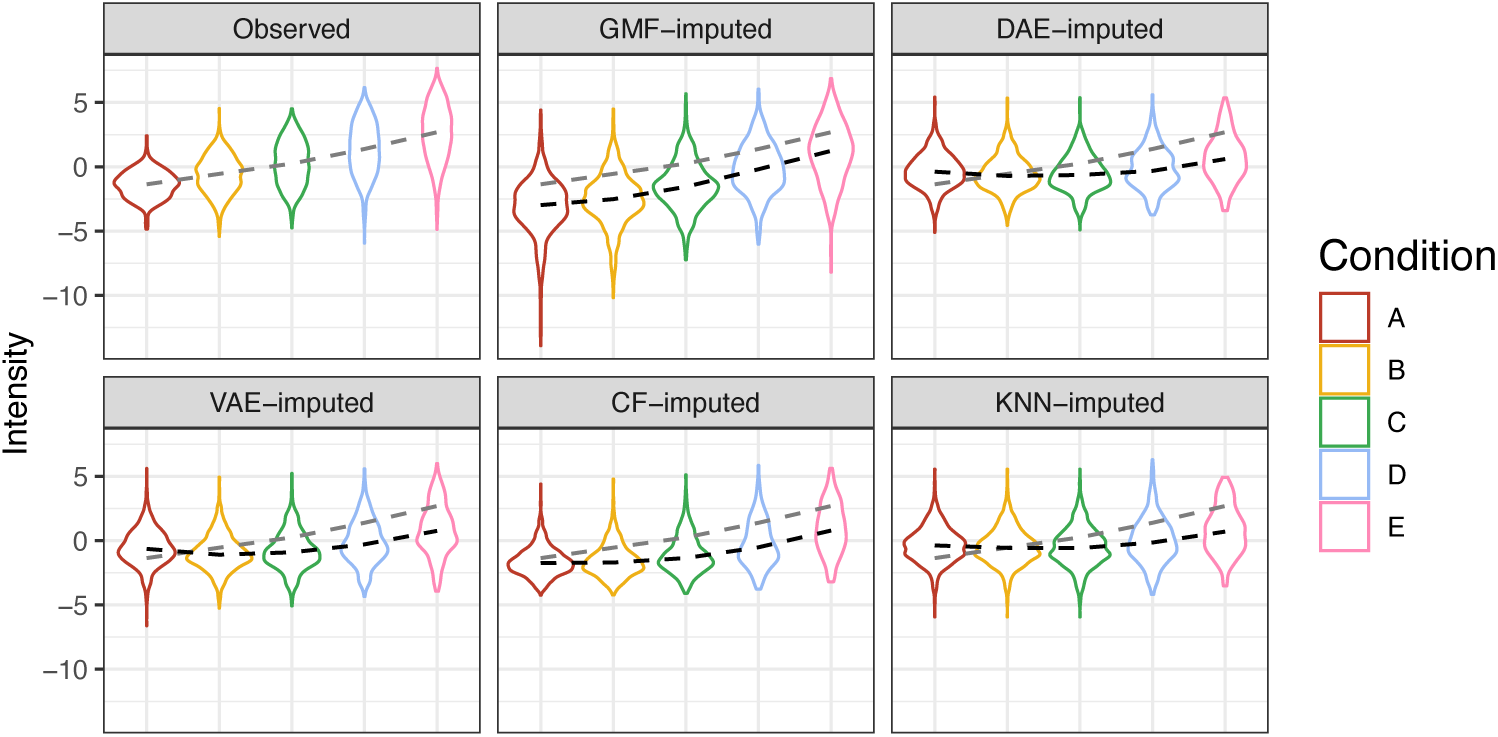
omicsGMF imputation accounts for missingness due to low abundance. Distributions of peptide intensities from human spike-in proteins are shown in function of the spike-in condition for the CPTAC data excluding Lab 1 that suffered from ionization issues. The first panel shows the distribution of observed values, and the other panels show the distributions of imputed intensities by omicsGMF, DAE, VAE, CF and KNN imputation respectively. The grey dashed line represents the median of the observed values, and the black dashed line the median of the imputed values.

Data exploration upon dimensionality reduction in the bulk, label-free CPTAC spike-in study (Fig. 3, Panel C; Supplementary Fig. 7), shows that intensity profiles from the Orbitrap at site 86, here referred to as Lab 1, deviates from those of the other labs. Interestingly, it has been previously reported that ionization issues occurred in Lab 1 while analyzing the samples from conditions A-C [16]. This is also clear from the vast number of missing values that can be observed for this lab (Supplementary Fig. 8), which has been partially overcome for the acquisition of samples from spikein condition D-E. omicsGMF has superior visualization of the data. It captures the spike-in concentration as most important source of variability in its first latent variable, and that of the technical lab-to-lab variability in its second latent variable. Upon removing the data from Lab 1 from the analysis, better visualizations are obtained for all methods (Fig. 3, Panel D; Supplementary Fig. 9). However, omicsGMF is still the only method that can distinguish between the most challenging conditions A and condition B. Indeed, many missing values for the differentially abundant UPS PSMs occur in these conditions due to their low spike-in concentrations. PCA upon imputation with the state-of-the-art tools, however, does not provide a good separation between the lowest spike-in conditions, suggesting that the performance of the dimensionality reduction is affected by the imputation of the high proportion of missing values for the differentially abundant features.

The graph contrastive deep learning framework scPROTEIN [18], developed explicitly for single-cell proteomics data, failed to generate sensible embeddings for both the Petrosius and Leduc datasets (Supplementary Fig. 10). This could be due to errors in its initial training step, which estimates the quality of the signal of each cell. Despite following the (rather limited) documentation, these errors could not be resolved. Therefore, scPROTEIN is not considered in the remainder of this manuscript.

The competing workflows also do not infer or provide guidelines on the number of latent factors for imputation nor for downstream applications, such as clustering, UMAP and t-SNE visualizations, or for correction for unknown batch effects and confounders, amongst others. Interestingly, omicsGMF allows selection of the optimal number of latent factors and/or the inclusion for known covariates by using cross-validation, information criteria, such as AIC and BIC, or scree plots, all of which can guide the user in different downstream applications.

Fig. 4 shows results for omicsGMF-based selection of the number of relevant latent variables and known covariates with cross-validation. Cross-validation on the Petrosius study (Fig. 4, panel A) shows that the optimal number of latent variables is five. As expected, when we incorporate the inhibitor treatment in omicsGMF, stem cells of both treatments nicely overlap in one homogeneous cluster when visualized in the first two latent variables (Supplementary Fig. 11). Remarkably, the optimal number of latent variables then reduces to four, which shows that the inclusion of one additional categorical dummy variable replaces one latent variable for optimal data representation. Note that the inhibitor treatment is also perfectly confounded with the acquisition time, so we cannot rule out that the difference we observe between the two populations of cells in the plot without known covariates is due to technical effects (Fig. 3, panel A). Nevertheless, the application clearly shows the strengths and relevance of omicsGMF in practice: omicsGMF captures this variability better than the existing methods in an unsupervised analysis and can effectively remove it from the analysis for applications that aim to cluster cells upon removal of the treatment or batch effect.

For the Leduc study, around forty latent variables are needed when no known covariates are fed to omicsGMF (Fig. 4, panel B). However, the cross-validation error dramatically reduces when correcting for the TMT run effect and omicsGMF then only needs ten latent variables. These results clearly indicate the importance of accounting for the run-to-run variability in this study, which is difficult to pick up with latent variables as the experiment involves many TMT runs with relatively few single cells profiled per run.

The results for the CPTAC study are particularly insightful (Fig. 4, panel C). Indeed, cross-validation for an omicsGMF analysis conducted without incorporation of known covariates identifies six latent variables. However, the introduction of the “Lab” variable in the omicsGMF model, produces a cross-validation score that is on par, albeit with only four latent variables: a reduction which matches with the inclusion of two categorical dummy variables representing Lab 2 and Lab 3, as Lab 1 serves as the reference group. Upon incorporating an additional categorical variable for spike-in condition, the optimal number of latent variables is further reduced to three. Interestingly, omicsGMF that correct for the lab variable seems to capture the variability of the spike-in condition with a single additional latent variable, although incorporating the spike-in condition explicitly in the omicsGMF model included four additional categorical dummy variables. This suggests that omicsGMF detects the linear increase in

UPS protein levels across conditions using a single, continuous, latent variable, which aligns with the actual experimental design. However, the cross-validation performance that includes the spike-in concentration is clearly inferior compared to omicsGMF models that exclude known covariates or account only for a batch effect associated with lab, probably due to more generous model with a factor for spike-in concentration overfitting the data.

### Imputation of missing values

The final mean estimates obtained from omicsGMF can also be utilized for the imputation of missing values while accounting for latent factors, as well as for known sample- and feature-level covariates. To evaluate imputation performance, we simulate missingness both completely at random (MCAR) and missing not at random (MNAR) due to low abundance (see Methods). The mean absolute error (MAE) between the imputed and original values is then assessed across ten different random seeds, considering scenarios where 25%, 50%, and 75% of the masked values are attributed to low abundance.

On the label-free Petrosius dataset (Supplementary Fig. 12 and 13), omicsGMF demonstrates superior imputation performance compared to PIMMS neural networkbased approaches DAE, VAE and CF, as well as to traditional methods such as NIPALS, KNN, QRILC, and zero or minimum imputation. Notably, omicsGMF outperforms other methods, particularly for MNAR. Despite being specifically designed for MNAR imputation, QRILC, zero, and minimum imputation result in dramatically higher error rates for MNAR data. For MCAR, however, methods that account for the covariance structure of the dataset i.e., omicsGMF, DAE, VAE, CF, and NIPALS, demonstrate comparable performance.

On the TMT-labeled Leduc dataset (Supplementary Fig. 14 and 15), omicsGMF accounting for batch effects, demonstrates an improved performance compared to all other imputation methods. This finding suggests that when complex technical artifacts are present, effective imputation of missing values requires explicit adjustment for these effects, both for MCAR and MNAR data. Although CF, DAE, and VAE were not designed for TMT-labeled data, they nonetheless outperform conventional imputation methods KNN, QRILC, zero, and minimum imputation.

For the label-free CPTAC dataset, omicsGMF achieves substantially lower imputation errors across the entire CPTAC study (Supplementary Fig. 16 and 17) as well as when only considering the subset excluding data from Lab 1, which is known to suffer from ionization issues (Supplementary Fig. 18 and 19). Notably, even when using the full CPTAC dataset and calculating the imputation performance exclusively for Lab 1, omicsGMF outperforms competing methods (Supplementary Fig. 20 and 21). This result suggests that omicsGMF not only provides more accurate imputation, but also effectively captures the underlying structure of the data.

omicsGMF has the advantage that it allows for the selection of the optimal number of latent factors prior to imputation. For a fair comparison we also ran DAE, VAE and CF with the number of latent factors selected by omicsGMF (Supplementary Fig. 22). However, this did not consistently improve the performance of these methods, indicating that those methods require their own hyperparameter optimalization.

Interestingly, the CPTAC study, with known ground truth for differentially abundant proteins, also allows the evaluation of the imputations for missing values in the original dataset i.e., without masking values. Fig. 5 presents the distribution of observed values for spike-in proteins, stratified by spike-in condition, alongside those generated by different imputation methods. This analysis excludes data from Lab 1, which is affected by ionization issues. Notably, omicsGMF successfully captures the concentration gradient in the missing values for UPS proteins, whereas other methods fail to recover this trend, particularly in the low spike-in conditions A, B, and C. Additionally, omicsGMF systematically imputes missing values lower than the observed ones, suggesting that missingness is driven by low abundance combined with lower ionization efficiency for specific peptide species. In contrast, other methods tend to shift imputed values toward higher intensities, sometimes exceeding those observed in conditions A, B, and even C, indicating difficulties in accurately imputing MNAR data. For non-differentially abundant yeast proteins, the imputed value distributions remain similar across conditions but are consistently lower than their observed counterparts (Supplementary Fig. 23). Interestingly, in condition E, the omicsGMF imputations are further shifted downward, mirroring a trend also seen in the observed yeast protein intensities. This is likely due to increased ionization competition caused by peptide overspiking in this condition, which omicsGMF seems to pick up on.

When including the Orbitrap data from Lab 1, which suffers from ionization issues, the omicsGMF imputation distributions for missing UPS peptides remains consistent for Labs 2 and 3, but deviates for Lab 1 (Supplementary Fig. 24). Specifically, omicsGMF imputes higher intensities in conditions A, B, and C, while imputations for conditions D and E are lower, reflecting the adjustments made to correct the ionization issue in Lab 1. However, the relative differences between conditions A, B, and C, and those between D and E remain preserved. Again, omicsGMF appears to better account for ion competition effects in yeast PSMs at higher UPS spike-in conditions. Hence, both the results for masked values in the simulation study and the results for actual missing values in the original CPTAC data indicate that omicsGMF better captures the real structure in the data than its competitors.

### Differential analysis

The CPTAC experiment provides a ground truth for differential abundance (DA) analysis, where 48 human UPS DA proteins are spiked at different concentrations alongside 1,477 non-DA yeast proteins. DA was inferred using msqrob2 [25, 26], incorporating a main effect for treatment and a block effect for lab, both on the original non-imputed data and on data processed with different imputation methods.

We first focus on the analysis using data from Labs 2 and 3. Fig. 6, panel A shows that only the workflows using non-imputed data and omicsGMF-imputed data provide reliable inference between the two lowest spike-in conditions, A and B. Other imputation methods generate numerous false positives in their top-ranked results. This outcome aligns with previous observations in Fig. 5, where most methods impute similar values for missing UPS peptides across these conditions. For comparisons between B vs. C and A vs. C, CF’s performance improves considerably, but still falls short of omicsGMF. Meanwhile, other state-of-the-art imputation methods continue to struggle with reliable DA inference. As the comparisons shift to higher spike-in concentrations, the performance of all methods improves, but omicsGMF consistently outperforms the existing methods (Supplementary Fig. 25).

**Fig. 6.**
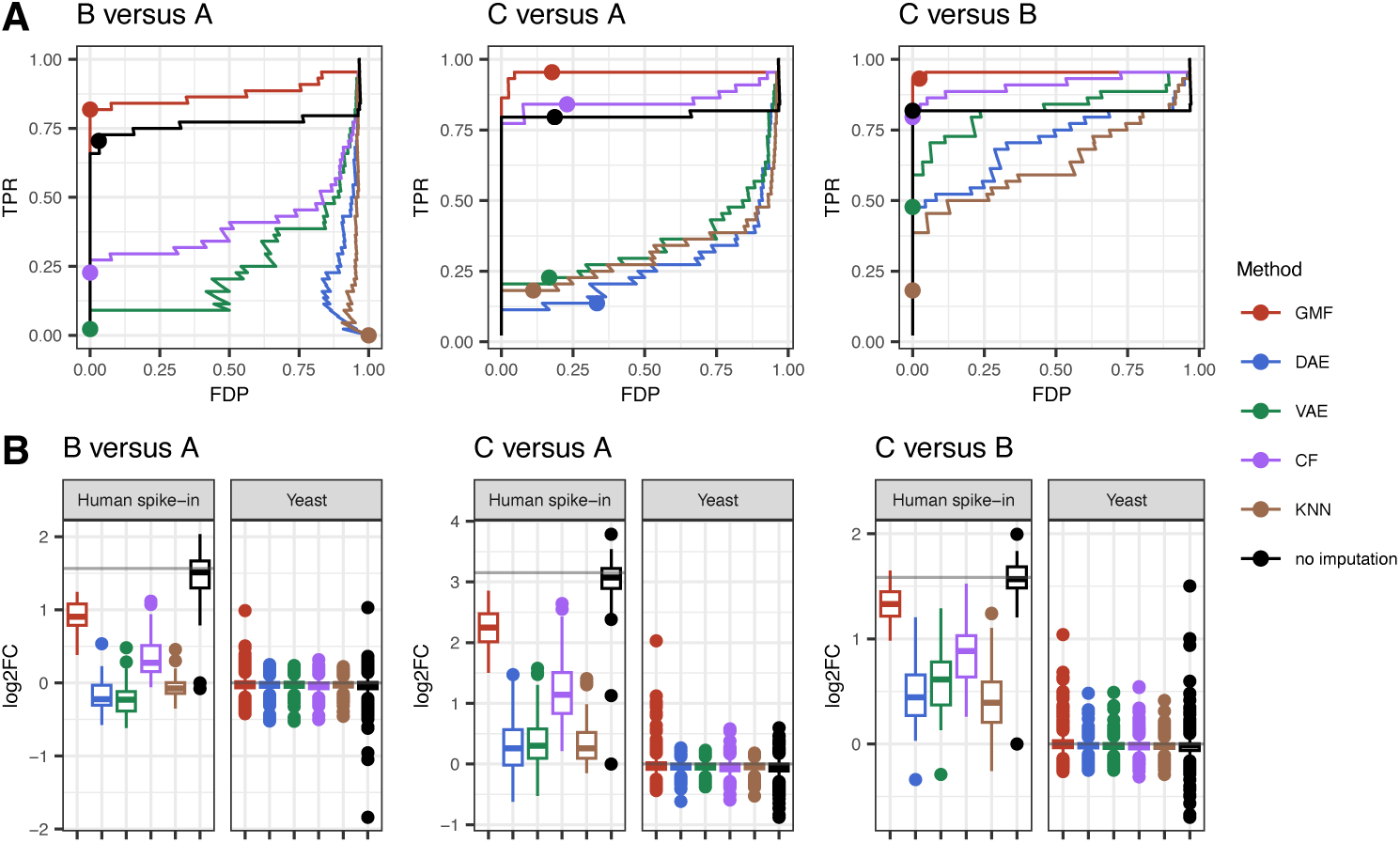
omicsGMF imputation leads to better downstream differential abundance analysis. Performance evaluation of differential abundance analyses using msqrob2 [25, 26] on the CPTAC dataset [16]. Results for the comparisons between the lowest spike-in concentrations B versus A, C versus A and C versus B are shown. Data from Lab 1 are excluded due to ionization issues. Human UPS proteins are differentially spiked between the conditions, with yeast background proteins as true negative control. Panel A shows the true positive rate (TPR) in function of the false discovery proportion (FDP). The dots on each curve represent working points when the FDR level is set at the nominal 5% level. Panel B shows the estimated log2 fold changes (FC) by msqrob2 for both human spike-in proteins, and for reference yeast proteins. The grey line indicates the known log2 FC.

Fig. 6, Panel B, and Supplementary Fig. 26 show that all imputation methods result in an underestimation of the fold change for spike-in UPS proteins. However, omicsGMF is the only approach that produces fold change estimates reasonably close to the ground truth, confirming that other methods struggle to correctly impute missing values due to low abundance. Interestingly, the bias in fold change estimates for spike-in UPS proteins disappears when only the observed PSM intensities are used in the msqrob2 analysis. However, this comes at the cost of reduced sensitivity as seen in Panel A of Fig. 6, because fewer data points are available for many spike-in proteins due to missingness.

When incorporating data from Lab 1, which experienced ionization issues, omicsGMF still outperforms other methods for the B versus A and C versus A comparisons (Supplementary Fig. 27). However, its performance deteriorates for comparisons involving spike-in conditions D and E. This drop is due to omicsGMF accounting for the ionization issues in Lab 1 samples from conditions A-C, leading to a upward shift in the imputed missing values as compared to those from conditions D and E in Lab 1. Despite this shift, however, the imputation preserved the relative differences among conditions A-C and D-E. Interestingly, even in comparisons involving conditions D-E, omicsGMF’s fold change estimates remain significantly less biased than those of competing methods (Supplementary Fig. 28).

When reanalyzing the complete CPTAC data with an additional dummy variable for conditions D-E of Lab 1 in the msqrob2 analysis, thus accounting for the ionization issues flagged during QC, omicsGMF once again emerged as the top performer (Supplementary Fig. 29). Note that the results for other imputation methods also improved. Interestingly, this adjustment reduced the bias in fold change estimates for all comparisons involving conditions D and E (Supplementary Fig. 30). This highlights the importance of properly accounting for technical artifacts in the downstream DA analysis. It also underscores the critical role of thorough quality control, for which low-dimensional visualization with omicsGMF proved particularly insightful, as it was the only method that clearly captured all leading sources of variability, namely: the spike-in condition, the lab effect, and the impact of poor ionization.

We repeated this benchmark using another dataset, in which an *E. coli* proteome was mixed at varying concentrations within a constant human background [27]. Consistent with our previous findings, the performance of DA following omicsGMF imputation surpasses that of other imputation strategies, particularly at low spike-in concentrations (Supplementary Fig. 31, 32). Furthermore, the imputed values generated by omicsGMF again exhibit a closer alignment with the trends across conditions for quantified yeast and human PSMs than those imputed with the existing methods (Supplementary Fig. 33). These results reaffirm that omicsGMF imputation leads to more powerful downstream DA analyses than existing imputation strategies.

## Discussion

omicsGMF is a scalable, flexible, and generic tool that streamlines and enhances data processing and dimensionality reduction for bulk and single-cell proteomics. It retrieves unknown sources of variation while simultaneously addressing the massive proportion of missing values and the strong batch effects, which are characteristic of large-scale MS experiments. omicsGMF shows several advantages over its current state-of-the-art competitors.

First, omicsGMF improves the quality of dimensionality reduction. For instance, for the Petrosius data set [4], omicsGMF returns a homogeneous cluster of stem cells after correcting for the effect of treatment. We also showed that correcting for the run effect for multiplexed cells was key to uncover a previously reported subpopulation of melanoma cells in the TMT-labeled SCP data from [3]. Moreover, ignoring known sources of variation to retrieve them as latent variables is a useful feature for data exploration and QC. For instance, on the CPTAC data, we convincingly showed that omicsGMF best captured the different sources of variability in the experiment, while flagging outlying samples suffering from ionization issues.

Second, correctly specifying the number of reduced dimensions has an important impact on the quality of the dimensionality reduction and imputation, and hence on the downstream analysis steps, such as UMAP and t-SNE visualization, cell clustering, trajectory inference, differential abundance analysis, among others. omicsGMF offers an automated model selection approach to determine the optimal number of latent variables, in contrast to state-of-the-art workflows that rely on the user-defined parameters without providing clear guidelines.

Third, omicsGMF provides sensible and accurate imputation and consistently ranks as a top performer across different datasets, for both MCAR and MNAR. The improved imputation performance systematically resulted in a more sensitive differential abundance analysis. Moreover, we found that omicsGMF is the only method that imputes values in line with the expected concentration for spike-in proteins, following a similar trend to the observed intensities for those proteins.

Fourth, omicsGMF relies on matrix factorization with interpretable model parameters. In contrast to neural networks which optimize weights that bear no meaning regarding the experimental context, the parameters estimated by omicsGMF are directly associated with experimental attributes (Fig. 2). This interpretation feature does not come at the expense of performance since omicsGMF outmatches the PIMMS deep-learning framework, both on small and large-scale datasets. These findings provide compelling evidence that Gaussian linear models are relevant for proteomics data analysis. A similar observation was reported by [28], who compared simple linear models to deep-learning frameworks specifically designed for predicting gene perturbations in the context of single-cell RNA sequencing.

Fifth, omicsGMF offers a framework for dimensionality reduction that simultaneously tackles the challenges of batch effects and missing values. Up to now, these tasks were performed in multi-step workflows, but designing multi-step workflows is cumbersome. It requires expert knowledge about the different methods available for each step and requires the optimization of the sequence of steps and their parameters [15]. The optimization is difficult and time-consuming to validate objectively, refraining the experimental researcher to confidently explore their data. Moreover, each dataset is unique in terms of technical and biological effects, amount of missing values, type of missingness, hence requiring multi-step workflows to be re-optimized, a costly procedure which is often neglected. Finally, multi-step workflows attempt to solve data challenges that are intertwined since missingness is influenced by batch effects [5], so there is no guarantee such workflows can lead to optimal and reliable results. In contrast, omicsGMF provides an extremely valuable alternative that integrates all steps in a single model, for which the parameters are simultaneously estimated. This improves performance, robustness across datasets, and overall user experience.

Finally, omicsGMF is available as an open source R-package, providing a user-friendly interface and vignettes for omics applications through https://github.com/statOmics/omicsGMF, which will soon be available on Bioconductor. In conclusion, omicsGMF offers an off-the-shelf solution that empowers researchers to thoroughly explore and process small- and large-scale (single-cell) proteomics data, alleviating the need to invest time and effort in data analysis technicalities.

## Methods

### omicsGMF Gaussian matrix factorization model

Let *y_ij_* be the intensity of peptide *j* (*j* = 1*, …, J*) in sample *i* (*i* = 1*, …, n*), which we model as a random variable following a Gaussian distribution with mean ***µ****_ij_* :

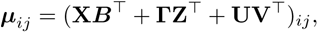

with **X** a *n* × *p* matrix with known sample-level covariates, such as batch or cell type or spike-in concentration, and ***B*** a matrix with its corresponding *J* × *p* regression parameters, **Z** a *J* × *q* matrix with known feature specific covariates, and ***L*** a matrix with its corresponding *n*×*q* regression parameters. Both **X** and **Z** can contain a column of ones, corresponding to gene-specific and sample-specific intercept, respectively. The *n* × *d* latent covariate matrix **U** and its *J* × *d* loading matrix **V** explain the residual variation that are not captured by the known covariates.

Because **U** and **V** are both unknown, these are estimated using an iterative process, i.e. with a block-wise stochastic gradient descent quasi-Newton method. In this manuscript, the core ideas of initialization and iterative estimation of the parameters are explained with a special focus on the huge missingness, which is characteristic for large-scale proteomics (LSP) and single cell proteomics (SCP) data. For further details upon block-wise parameter estimation, the adapting learning rate, and the smoothing of gradient and Hessian matrices, we refer the reader to our technical manuscript on the sgdGMF framework [19].

### Initialization of the parameters

Because parameter estimation cannot be done when the input matrix contains missing values, omicsGMF uses soft-imputation for the initialization and optimization of the parameters. This soft-imputation avoids using fixed values for the missing values, by updating their estimates in each iteration.

We adopt the notation of [29] and [30] and define the *n* × *J* matrix *P*_Ω_(**Y**),

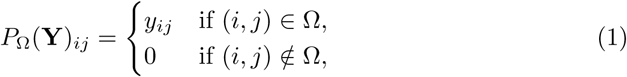

which is the projection of the *n* × *J* intensity matrix **Y** on the observed entries, and _Ω_ is the index set of the observed data. The complementary projection 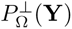 is defined via 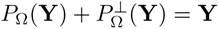.

Let ***µ***_0_ be the *n* × *J* matrix with feature means ***µ***_0*,ij*_ = mean(*P*_Ω_(**Y***_j_*)) used for initialization of the *n* × *J* imputed intensity matrix **Ŷ** :

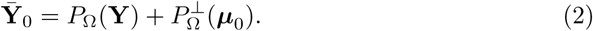

Using this imputed intensity matrix, the parameters related to the known sample- and feature-level covariates can be initialized. This can be achieved by maximum likelihood estimation or by minimizing the residual sum of squared errors:

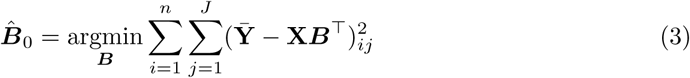

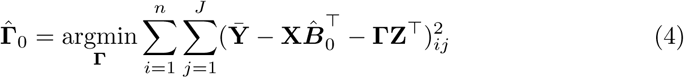

Then, we can initialize the latent covariate matrix **U** and its loadings **V** by performing a principal component analysis on the *n* × *J* working residuals matrix **E**:

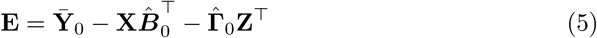

This gives an initial estimate of all the parameters ***B***, **Z**, **U** and **V** from which omicsGMF proceeds with optimization of these parameters.

### Optimization of the parameters

The optimization process consist of three consecutive steps. We consider iteration *t*, and iterate till convergence:

1. The imputed values are updated based on the current mean intensity estimates:

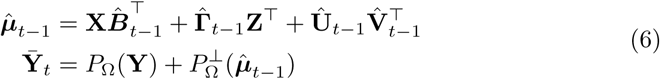

1. Update of ***B*** and **V** by minimizing the residual sum of squared errors:

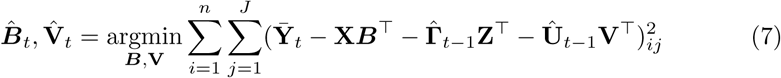

1. Update of ***L*** and **U** by minimizing the residual sum of squared errors:

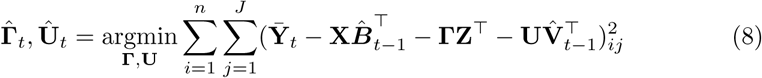

Upon convergence, **U** and **V** are orthogonalized via truncated singular value decomposition of **UV**^T^ in such a way that **U**^T^**U** is *d*×*d* diagonal matrix, with decreasing diagonal elements, and **V**^T^**V** is the *d* × *d* identity matrix, thus mimicking the parametrization conventionally used in PCA. Then, **U** can easily be used for visualization as they have the same interpretation as the scores in PCA. At convergence, the estimated means, ***µ***^*_t_*, can be used for imputation purposes.

Note that in steps 2 and 3, omicsGMF does not compute the optimal values of the parameters. Rather, it uses approximated updates via stochastic gradient descent with minibatch subsampling for computational efficiency, and only a few samples and features are considered in each update of the parameters. Therefore, such a local gradient refinement might not represent the optimal solution for the complete matrix. To mitigate this issue enhancing the algorithm’s stability while keeping a high level of computational efficiency, omicsGMF uses partial parameter updates and exponential gradient averaging. For more details, we refer the reader to the technical manuscript on the sgdGMF framework [19].

### Cross-validation to determine the number of latent covariates

To use omicsGMF, the number of latent variables included in the model, *d*, has to be selected. omicsGMF has the option to select these using the eigenvalues in a screeplot, information criteria, such as AIC or BIC, or cross-validation. In this manuscript, we consider 3-fold cross-validation to select *d*. During cross-validation, a subset of the observed data is masked as missing. Then, for various numbers of latent factors, omicsGMF imputes these values after convergence and calculates the residual sum of squared errors between the imputed and original values for the masked entries. The number of latent variables that minimizes the sum of squared errors is then considered the optimal number of latent variables and is used for the final fit without masking any observed values. The cross-validation results on the different datasets are available in Fig. 4.

### Benchmarking of imputation methods

To benchmark the different imputation methods, we consider the approach introduced by [11], which simulates both missingness completely at random (MCAR) and missingness due to low abundance, or missingness not at random (MNAR). In summary they consider:

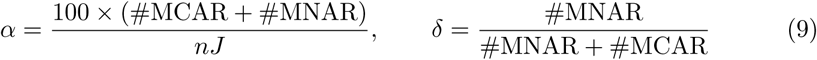

where *α* represents the percentage of observed values turned into missing values, and is here fixed at 10%. *δ* is the ratio of MNAR to the total missingness introduced in the dataset. This value is set at 25%, 50%, and 75%, to benchmark methods across a broad range of MNAR levels. This means that respectively 75%, 50%, and 25% of the missingness are MCAR. MNAR is based on a stochastic threshold. Assume a threshold matrix **W** from a Gaussian distribution parameterized by *µ_W_*= *c* and *σ_W_* = 0.01, where *c* is the *α^th^* quantile of the observed intensities. When the observed intensity for sample *i* and feature *j* is below its threshold **W***_ij_*, its value is censored by a Bernouilli draw with probably of censoring equal to *δα/*100. For more details, we refer to [11].

We conducted 10 simulations, each introducing different missing values for each MNAR percentage. For each simulation, the mean average error between the imputed and observed values were computed, for each entry that was masked during the simulation.

### State-of-the art tools for imputation in proteomics

We conduct an extensive comparison with the following bespoke methods from the proteomics literature for dimensionality reduction and imputation.

scPROTEIN [18] is a deep graph contrastive learning framework that searches for latent embeddings to visualize single-cell proteomics data. It consists of two stages. The first stage estimates the uncertainty of the peptide signals. However, this step led to errors in all datasets and, therefore, could not be used in our contribution. The second stage is contrastive learning to search the embedding of the samples. This tool can only be used for dimensionality reduction and not for imputation.

PIMMS DAE [13] is a denoising autoencoder that trains a deep neural network by masking values and attempting to reconstruct them. It is used to impute missing values and requires hyperparameters for the dimensionality of the latent space and the number of hidden layers. For the latter we used the default value of 512 hidden layers, while the dimensionality of the latent space was either set at the default value of 50 or at the value suggested by the cross-validation of omicsGMF.

PIMMS VAE [13] is a variational autoencoder similar to the denoising autoencoder but with a stochastic latent space optimized to follow a standard normal distribution. It is also used for imputation of missing values. The deep neural network is again trained by masking values and trying to reconstruct them. We again used the default value of 512 hidden layers, and have set the dimensionality of the latent space to either the default of 50 or at the one suggested upon cross-validation with omicsGMF.

PIMMS CF [13] is a collaborative filtering deep learning approach where both features and samples have trainable latent embedding spaces. The hyperparameter is the latent space of the separate sample and feature embeddings, and was either set to its default value of 30 or to the one suggested upon cross-validation with omicsGMF. KNN (K-nearest neighbours) [20] searches for *k* features (peptides or proteins) that are most similar to the one with missing values, using Euclidean distances. Each missing value is imputed by averaging the intensities of these features in its corresponding sample. Here, *k* is set to 10. We used NAguideR [31] as a wrapper for the impute package.

NIPALS (non-linear estimation by iterative partial least squares) [9] performs PCA by ignoring missing values when calculating the inner products. It does not require imputation to compute the matrix factorization. The dimensionality used corresponds to the one suggested upon cross-validation with omicsGMF. Further, NIPALS can also be used for imputation, by multiplying the scores and loadings of its matrix factorization. NIPALS cannot correct for known covariates.

QRILC [21] imputes missing values by drawing values from a truncated normal distribution estimated by quantile regression. This method was mainly developed to impute missing values based on missing due to low abundance. NAguideR [31] was used as a wrapper for the imputeLCMD package.

Finally, zero imputation and minimum imputation simply impute every missing values with 0 or the minimum value observed in the data, respectively, and are implemented in NAguideR [31].

Methods that can only impute data were followed by classical PCA to evaluate the dimensionality reduction upon imputation.

### Data

This manuscript builds its results from two single cell and two bulk proteomics datasets that were acquired with either TMT-labeling or label-free quantification.

First, the Petrosius dataset [4] originates from a label-free SCP experiment on mouse embryonic stem cells incubated in two distinct conditions i.e., a serum-free 2i condition (m2i) containing cytokine LIF with inhibitors for the MEK and GSK3 pathways, and a serum condition (m15) containing cytokine LIF, only. Data were acquired using an orbitrap Eclipse Tribrid mass spectrometer operated in data independent acquisition mode. The MS data were analyzed by the authors using Spectronaut v17 [32]. We performed quality control by removing cells with fewer than 750 detected peptides or a median log2-intensity lower than log2(7). Features with more than 90% missing values were also removed. The detected intensities were log2-transformed for variance-stabilization, and the intensities of each cell were centered with the median log2-intensity of the corresponding cell. The resulting dataset contains 525 cells with 4435 detected peptides, of which 58.5% of the values are missing.

Second, the Leduc dataset [3] was acquired using pSCoPE technology, which relies on TMT labeling and a prioritized data dependent acquisition strategy carried out by a Thermo Scientific Q-Exactive mass spectrometer. The MS data were quantified and identified using MaxQuant version 1.6.17 [33]. We performed quality control by filtering cells with fewer than 750 detected peptides or a median coefficient of variation greater than 0.5, calculated using the *medianCV perCell* function of the *scp* [15] package. Peptides with more than 90% missing values were removed, resulting in a dataset with 6280 peptides for 1508 cells and a total of 61.8% missingness. The intensities were log2-transformed for variance-stabilization, and the log2-intensities of each cell are centered using the median log2-intensity of the corresponding cell.

Third, data from an interlaboratory bulk label-free proteomics spike-in study of the CPTAC consortium was used [16]. Forty-eight human proteins were spiked in a background of yeast proteins at 5 different concentrations (*A* = 0.25*fmol/µl*, *B* = 0.76*fmol/µl*, *C* = 2.2*fmol/µl*, *D* = 6.7*fmol/µl* and *E* = 20*fmol/µl*) and three samples of each concentration were sent to three different labs resulting in data from forty-five different MS-runs. Hence, the ground truth on differential abundance is known, which is useful to assess the imputation of missing values for differential (UPS) and non-differential proteins (Yeast), and how the different imputation methods affects the downstream differential abundance analyses. The data were searched and quantified according to [25]. Again, peptides with more than 90% missing values were removed, resulting in a total of 10105 peptides remaining from 1477 yeast proteins and 44 human proteins and 42.5% missing values.

Finally, we used Shen dataset [27], another bulk proteomics label-free benchmarking effort consisting of E. coli proteins mixed in five different weight ratio’s in a background of the human reference proteome (in wt/wt percentage: a = 3%, b = 4.5%, c = 6%, d = 7.5 % and e = 9%). After normalization and filtering peptides with more than 90% missing values, the dataset consists of 28944 peptides from 756 E. coli proteins and 3954 human proteins with 18.8% of the values missing. The data originates from twenty MS-runs acquiring four samples from each spike-in condition.

## Data availability

All data used in this manuscript are publicly available. The Petrosius [4] and Leduc [3] datasets were downloaded using the scpdata [5, 15] package from Bioconductor. The unfiltered intensities of the CPTAC [16] and Shen data [27] are available through https://github.com/statOmics/GMFProteomicsPaper.

## Code availability

All code used to prepare and analyze the datasets, and produce the figures of this manuscript are available through https://github.com/statOmics/ GMFProteomicsPaper.

Finally, omicsGMF is published as an open source R-package through https://github.com/statOmics/omicsGMF, which will be available soon on Bioconductor. It builds on the sgdGMF framework, which is available through CRAN, https://CRAN. R-project.org/package=sgdGMF.

**Supplementary information.** Supplementary Information contains Supplementary Figures 1-33.

## Ethics declarations

