## Supplementary Information for "omicsGMF: a multi-tool for dimensionality reduction, batch correction and imputation applied to bulk- and single cell proteomics data"

### Supplementary Figures

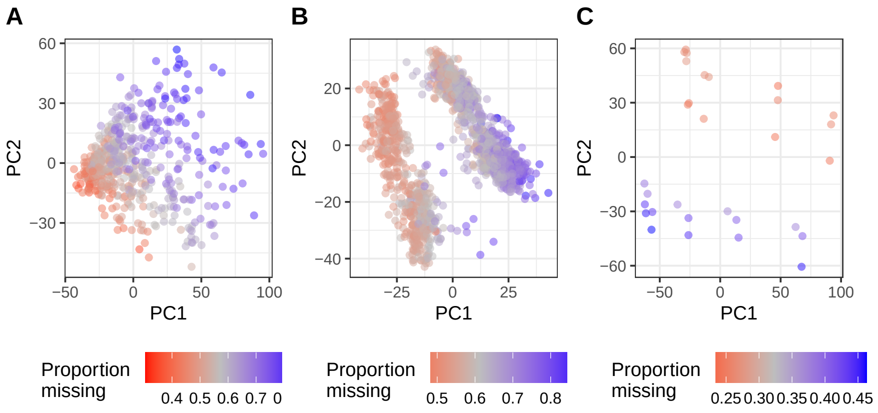


**Supp. Fig. 1** Panels A, B and C show data from the label-free single-cell Petrosius study [1], the labeled single cell Leduc dataset [2], and the data from the label-free bulk CPTAC spike-in study [3], respectively. Cells are coloured by their proportion of missing values. The principal components seem to be correlated with the proportion of missing values. All low dimensional visualizations were obtained with state-of-the-art CF-imputation [4] followed by PCA.

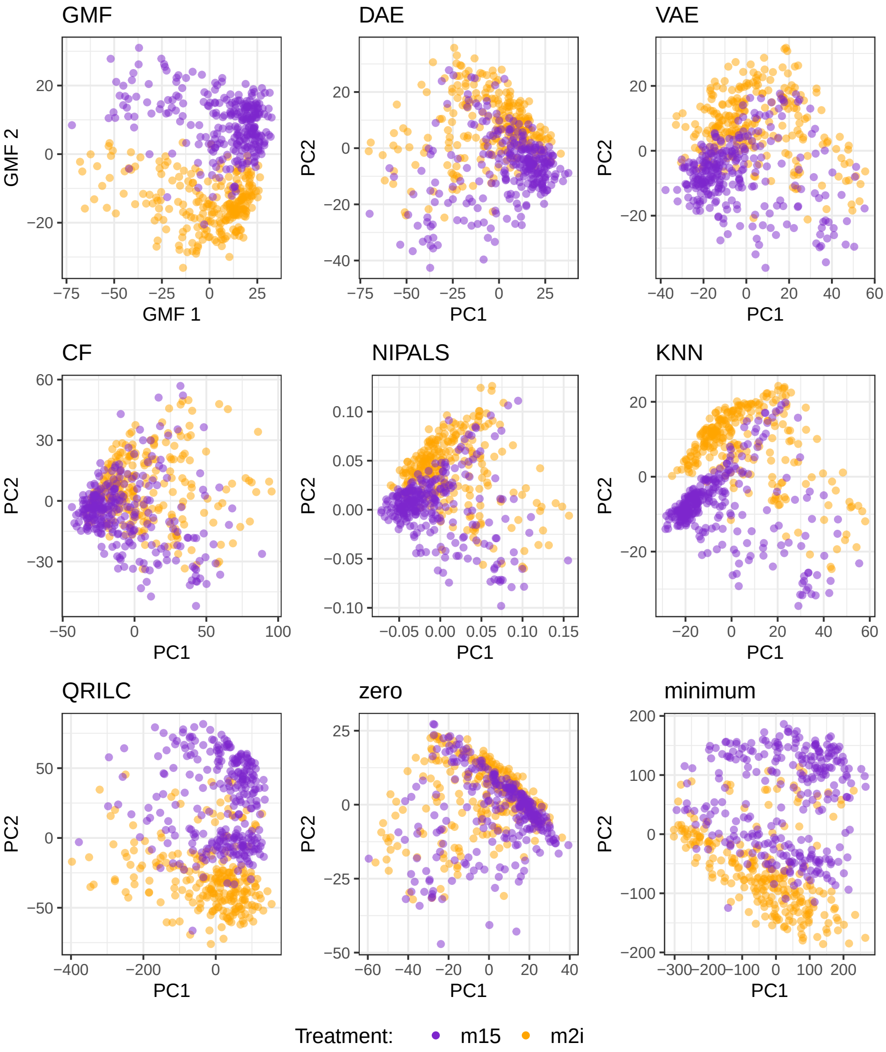


**Supp. Fig. 2** Low-dimensional visualization of the Petrosius data [1], colored by inhibitor treatment. omicsGMF and NIPALS directly estimate latent factors that have a similar interpretation as regular PCA. The other visualizations are obtained by imputation of missing values using DAE, VAE, CF, KNN, QRILC, zero and minimum, followed by PCA.

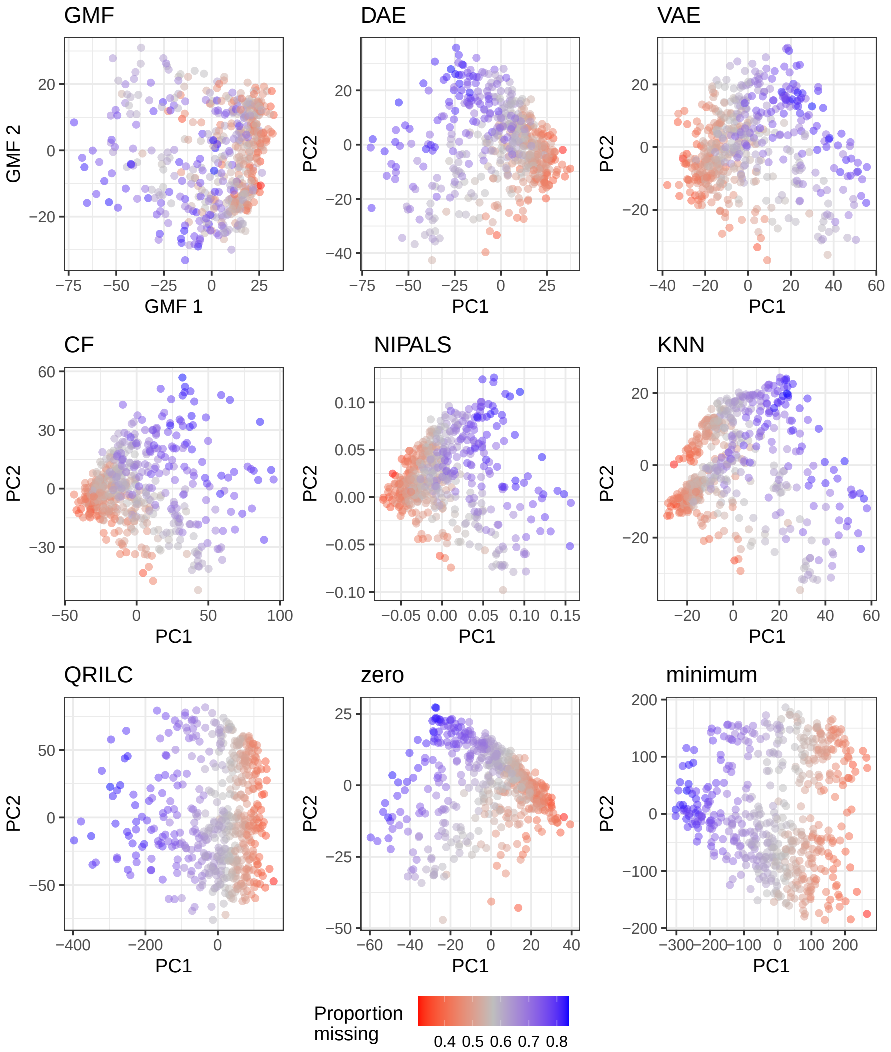


**Supp. Fig. 3** Low-dimensional visualization of the Petrosius data [1], colored by proportion of missing values in the cell. omicsGMF and NIPALS directly estimate latent factors that have a similar interpretation as regular PCA. The other visualizations are obtained by imputation of missing values using DAE, VAE, CF, KNN, QRILC, zero and minimum, followed by PCA.

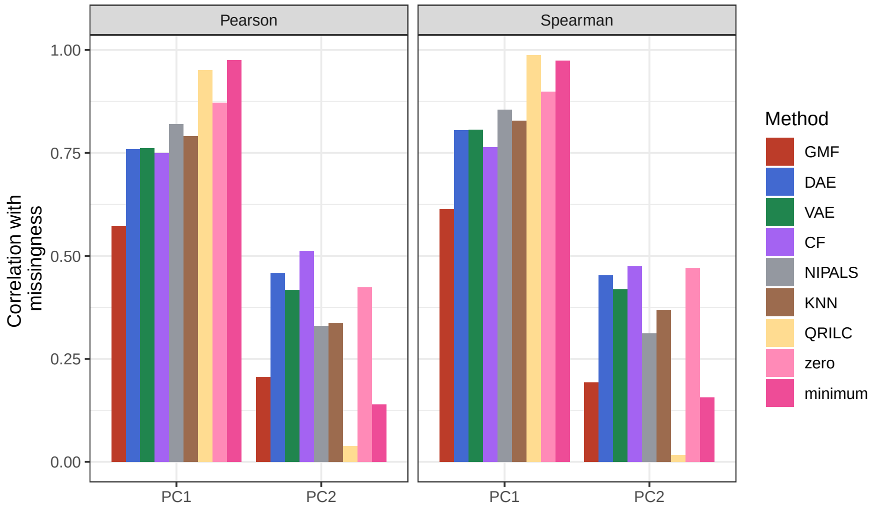


**Supp. Fig. 4** Barplots of the Pearson (left) and Spearman (right) correlations between the first two principal components and the proportion of missing values in a cell from the Petrosius dataset [1].

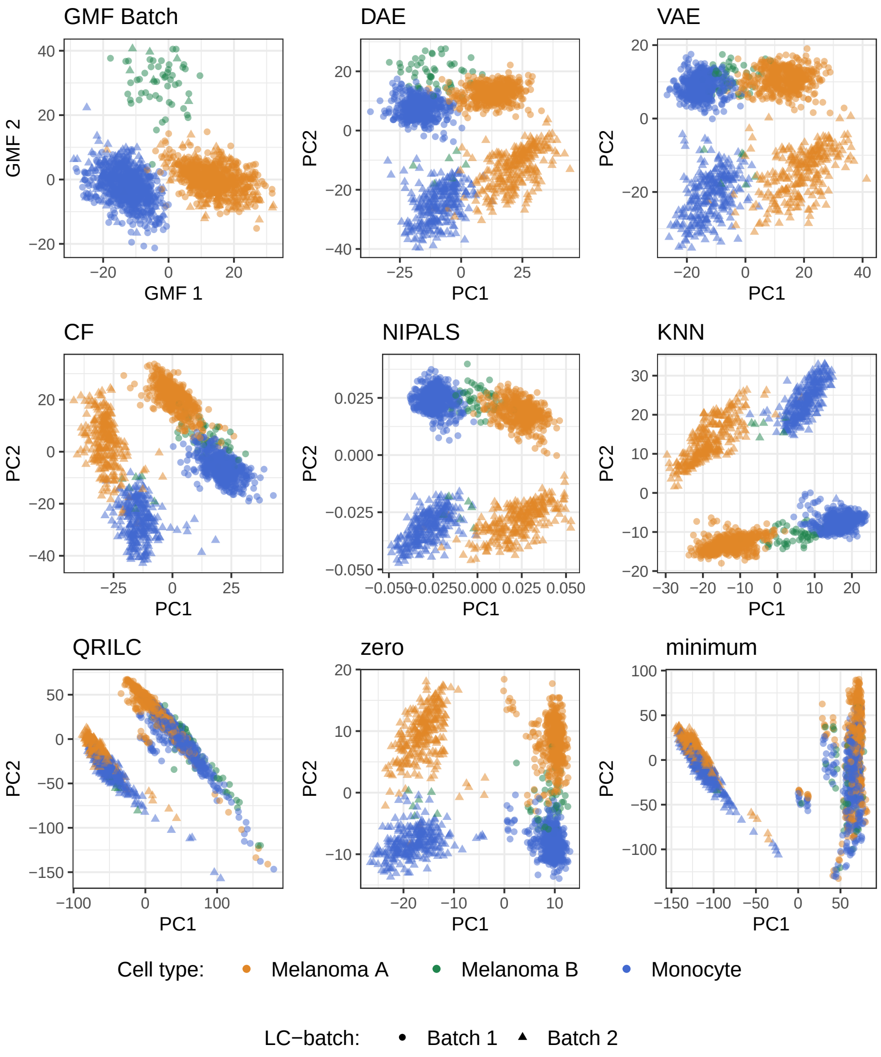


**Supp. Fig. 5** Low-dimensional visualization of the Leduc data [2], colored by cell type. omicsGMF and NIPALS directly estimate latent factors that have a similar interpretation as regular PCA. Here, omicsGMF directly accounts for known batch effects, resulting in a better representation of the biological signal. The other visualizations are obtained by imputation of missing values using DAE, VAE, CF, KNN, QRILC, zero and minimum, followed by PCA.

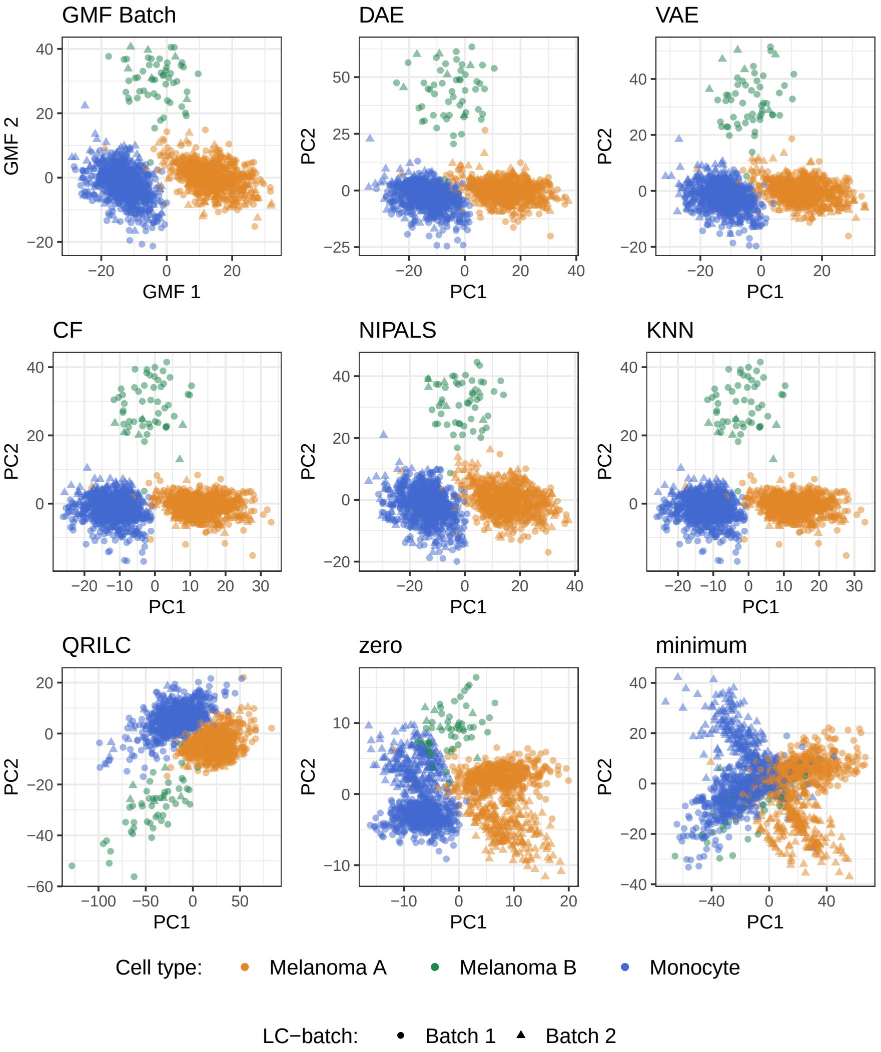


**Supp. Fig. 6** Low-dimensional visualization of the Leduc data [2] after batch-correction, colored by cell type. omicsGMF directly accounts for known batch effects. The other visualizations are obtained by imputation of missing values using DAE, VAE, CF, NIPALS, KNN, QRILC, zero and minimum, followed by batch-correction using linear regression and PCA on the remaining residuals.

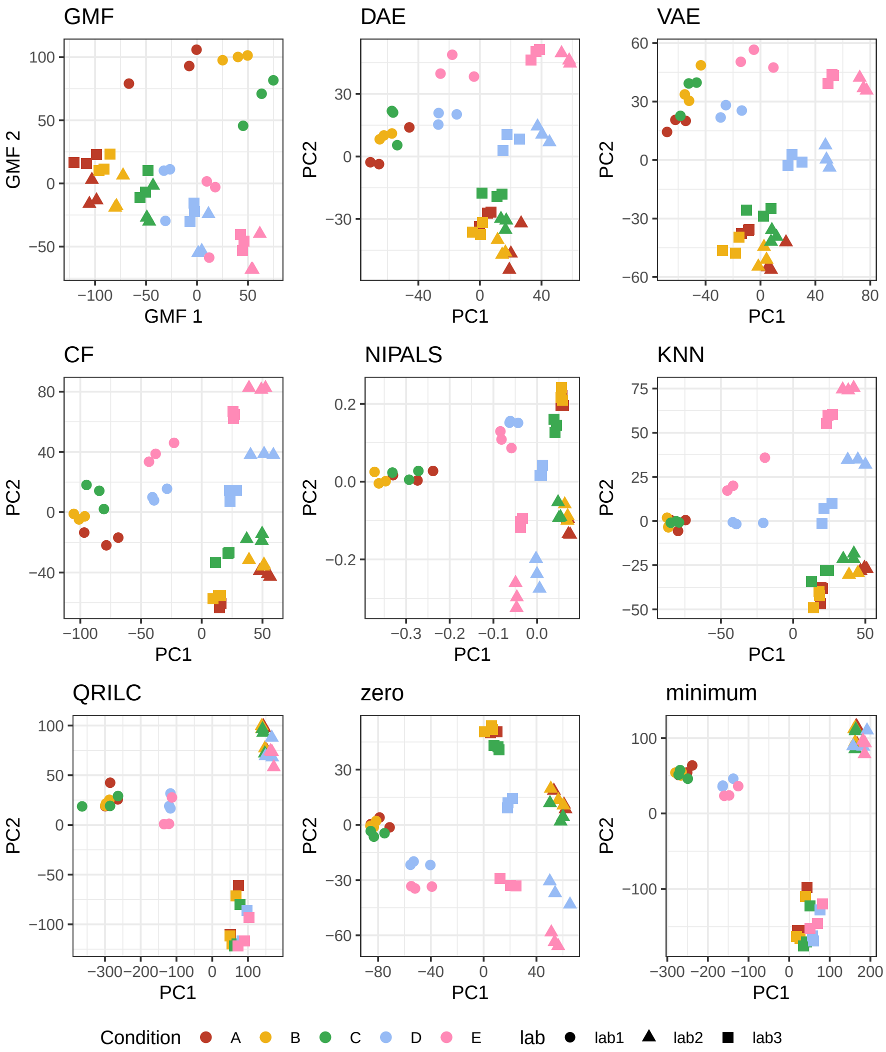


**Supp. Fig. 7** Low-dimensional visualization of the complete CPTAC data [3], colored by spike-in concentration of human proteins. Distinct marker shapes indicate the different labs. omicsGMF and NIPALS directly estimate latent factors that have a similar interpretation as regular PCA. The other visualizations are obtained by imputation of missing values using DAE, VAE, CF, KNN, QRILC, zero and minimum, followed by PCA.

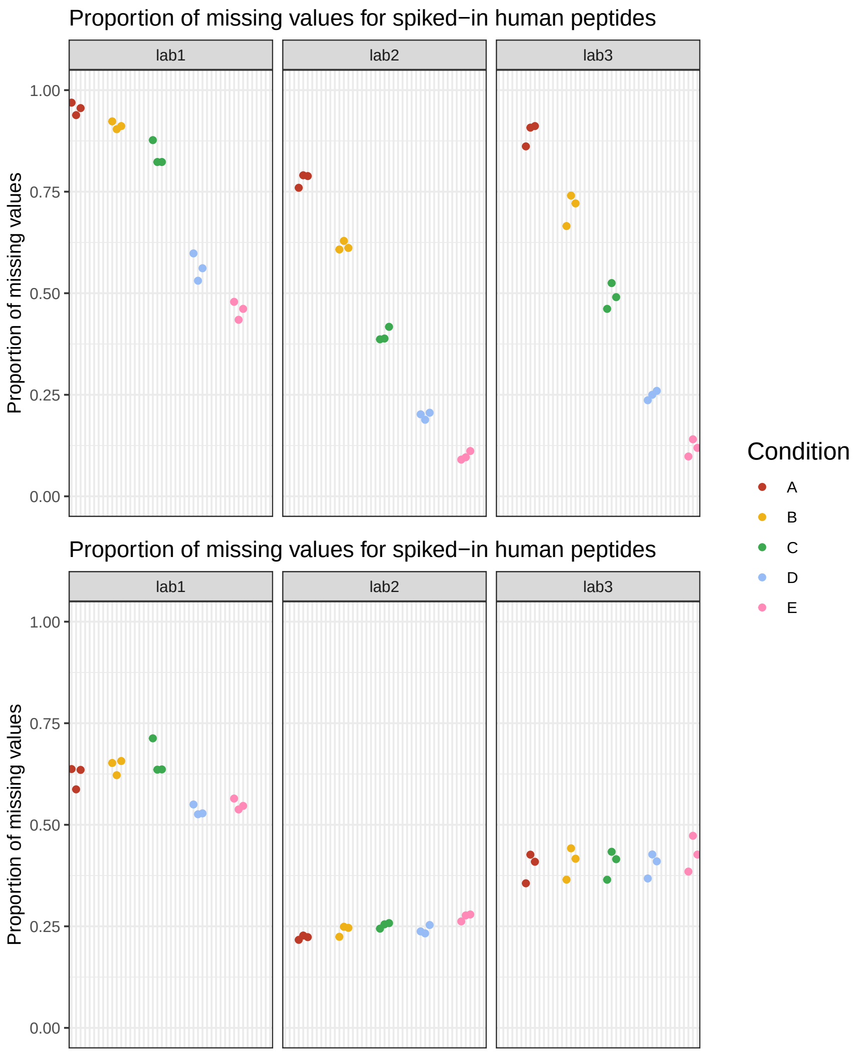


**Supp. Fig. 8** Proportion of missing values for each sample of the complete CPTAC dataset [3], for spike-in human peptides (top) and the background yeast peptides (bottom) respectively. The proportion of missing values is represented in function of the spike-in concentration, and is stratified per lab.

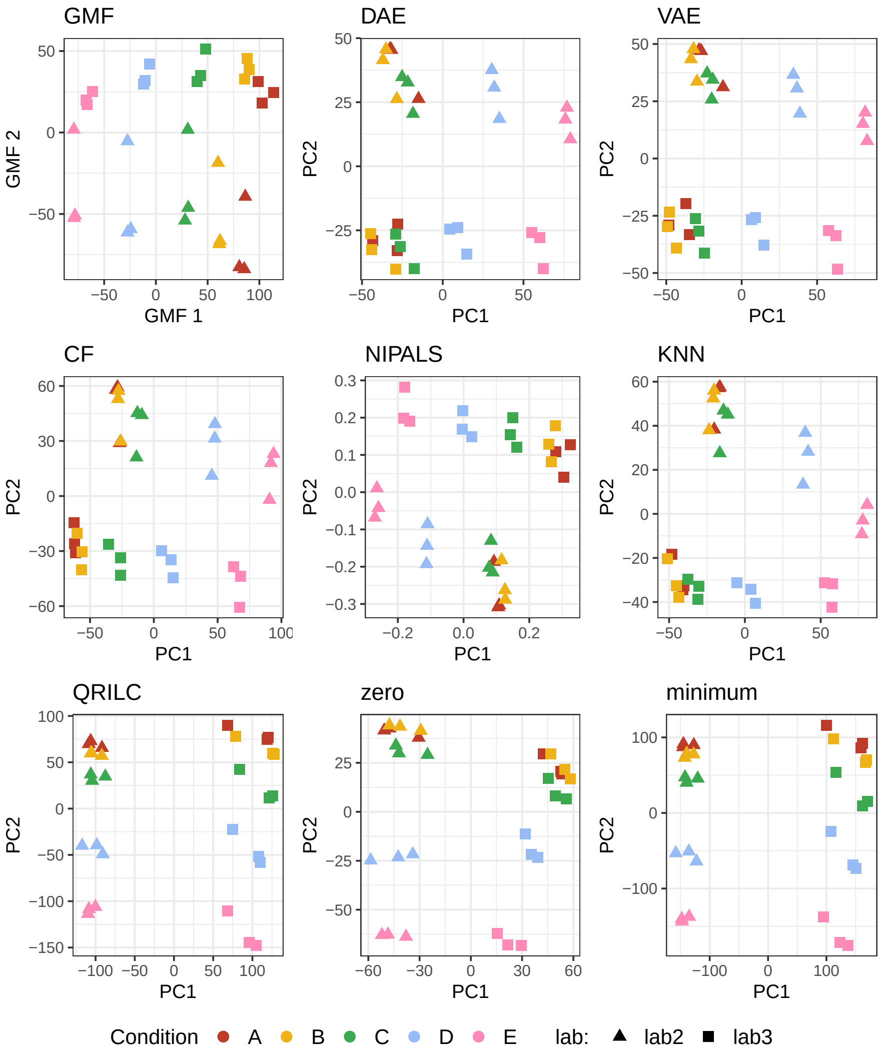


**Supp. Fig. 9** Low-dimensional visualization of the CPTAC data [3] upon excluding Lab 1, which was reported to suffer from ionization issues. Samples are colored by spike-in concentration of human proteins, and different labs are represented with a distinct marker shape. omicsGMF and NIPALS directly estimate latent factors that have a similar interpretation as regular PCA. The other visualizations are obtained by imputation of missing values using DAE, VAE, CF, KNN, QRILC, zero and minimum, followed by PCA.

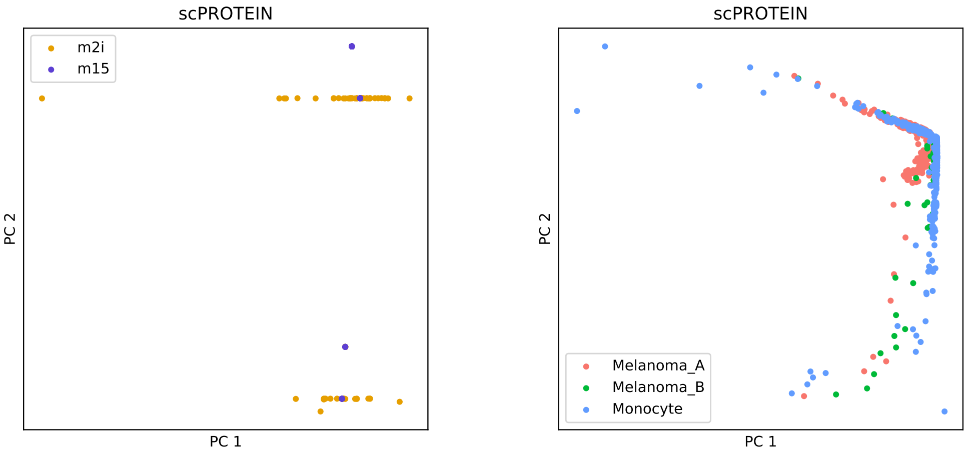


**Supp. Fig. 10** Low-dimensional visualization of the Petrosius [1] (left), and Leduc [2] (right) data by the scPROTEIN [5] workflow. Samples are coloured by inhibitor treatment and cell types respectively. No sensible embeddings are obtained by scPROTEIN, which could be due to errors in its initial training step estimating the quality of the signal of each cell.

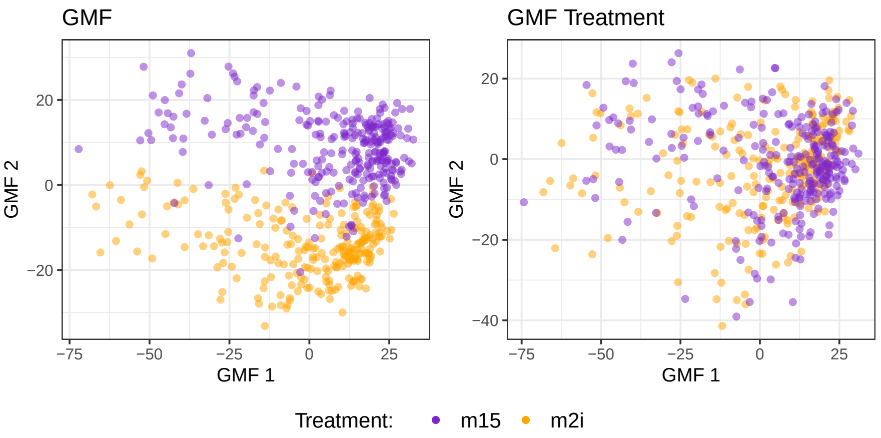


**Supp. Fig. 11** Low-dimensional visualization of the Petrosius data [1], colored by inhibitor treatment. omicsGMF is used both without (left) and with (right) a dummy variable for the inhibitor treatment. Clearly, the treatment effect is filtered out if this known covariate is accounted for, resulting in visualizations that control for this effect. This showcases the use of omicsGMF when differently treated cells have to be clustered together.

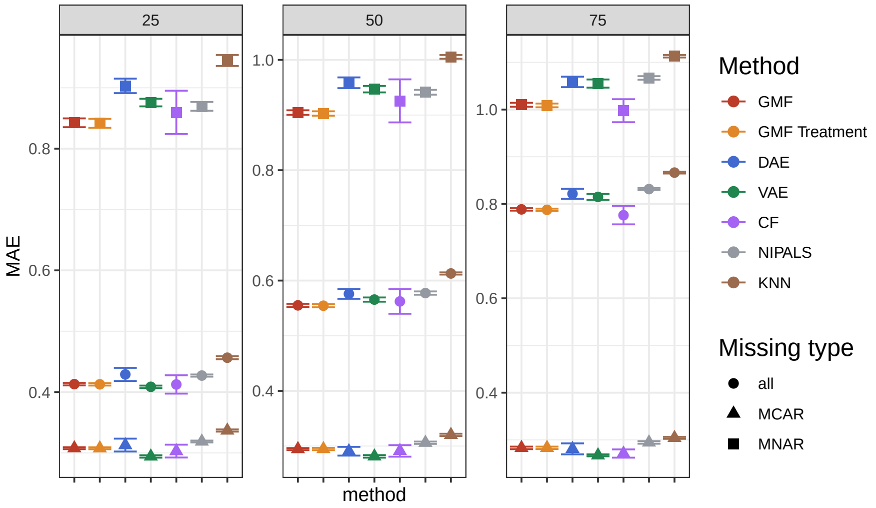


**Supp. Fig. 12** Mean average error (MAE) of the imputed values in the Petrosius dataset [1] evaluated for omicsGMF (GMF), omicsGMF including a dummy variable for the treatment effect (GMF Treatment), DAE, VAE, CF, NIPALS and KNN-imputation. Missing values were simulated according to the procedure described by [6] (see Methods), which introduces both missing completely at random (MCAR) and missing not at random (MNAR) values in predefined proportions. In this study, the proportions of MNAR masked values were set to 25% (left), 50% (middle), and 75% (right). For each condition, 10 different random seeds were used, and the mean MAE across these seeds is shown, with error bars representing the standard error of the MAE. The MAE was calculated exclusively for masked values based on the difference between the imputed values and the original observed values prior to masking. Distinct marker shapes indicate the MAE for only MCAR masked values, MNAR masked values, and for all masked values combined (all).

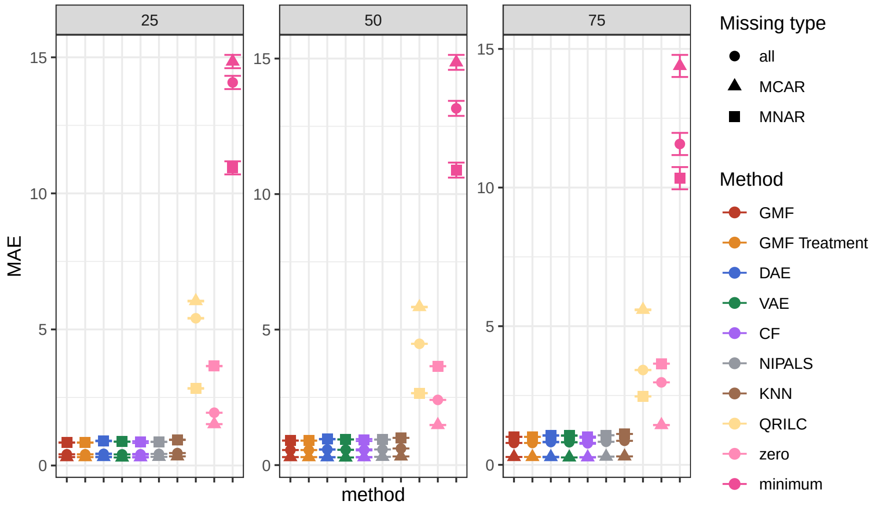


**Supp. Fig. 13** Same figure as Supp. Fig 12, but with additional methods. Mean average error (MAE) of the imputed values in the Petrosius dataset [1] evaluated for omicsGMF (GMF), omicsGMF including a dummy variable for the treatment effect (GMF Treatment), DAE, VAE, CF, NIPALS, KNN, QRILC, zero and minimum imputation. Missing values were simulated according to the procedure described by [6] (see Methods), which introduces both missing completely at random (MCAR) and missing not at random (MNAR) values in predefined proportions. In this study, the proportions of MNAR masked values were set to 25% (left), 50% (middle), and 75% (right). For each condition, 10 different random seeds were used, and the mean MAE across these seeds is shown, with error bars representing the standard error of the MAE. The MAE was calculated exclusively for masked values based on the difference between the imputed values and the original observed values prior to masking. Distinct marker shapes indicate the MAE for only MCAR masked values, MNAR masked values, and for all masked values combined (all).

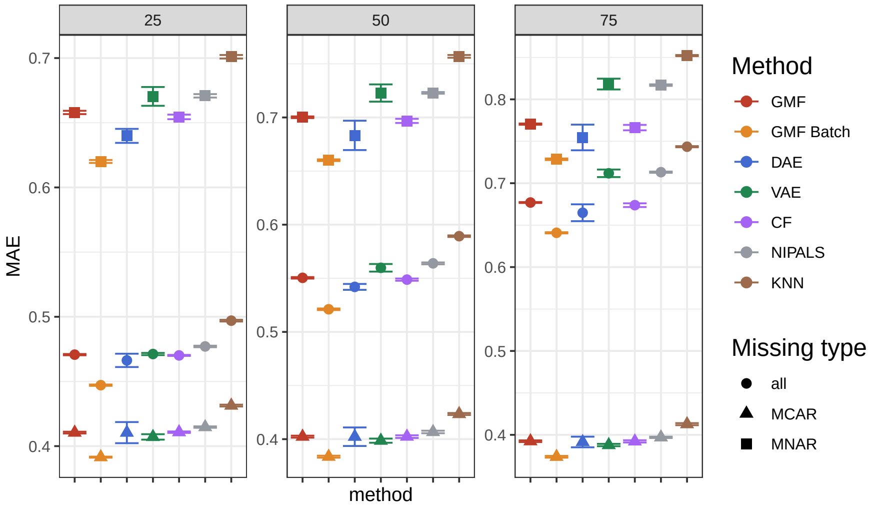


**Supp. Fig. 14** Mean average error (MAE) of the imputed values in the Leduc dataset [2] evaluated for omicsGMF (GMF), omicsGMF including 142 dummy variables for the batch effects (GMF Batch), DAE, VAE, CF, NIPALS and KNN-imputation. Missing values were simulated according to the procedure described by [6] (see Methods), which introduces both missing completely at random (MCAR) and missing not at random (MNAR) values in predefined proportions. In this study, the proportions of MNAR masked values were set to 25% (left), 50% (middle), and 75% (right). For each condition, 10 different random seeds were used, and the mean MAE across these seeds is shown, with error bars representing the standard error of the MAE. The MAE was calculated exclusively for masked values based on the difference between the imputed values and the original observed values prior to masking. Distinct marker shapes indicate the MAE for only MCAR masked values, MNAR masked values, and for all masked values combined (all).

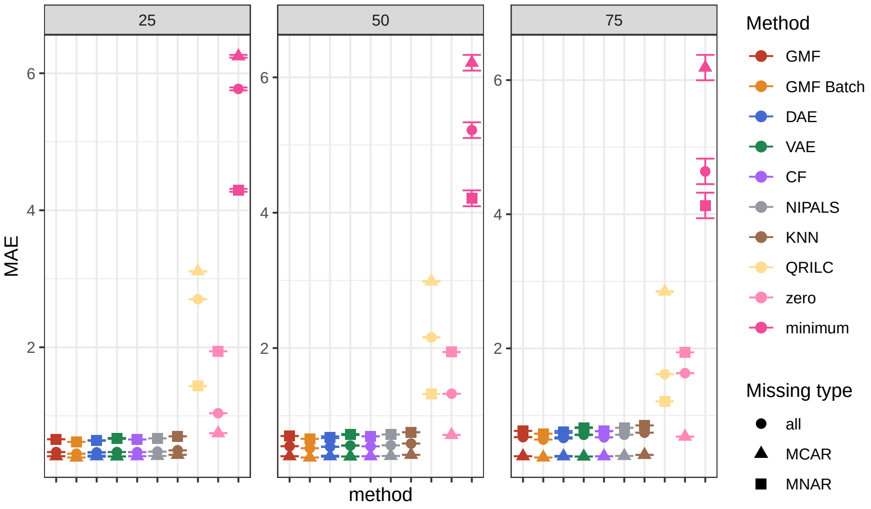


**Supp. Fig. 15** Same figure as Supp. Fig 14, but with additional methods. Mean average error (MAE) of the imputed values in the Leduc dataset [2] evaluated for omicsGMF (GMF), omicsGMF including 142 dummy variables for the batch effects (GMF Batch), DAE, VAE, CF, NIPALS and KNN, QRILC, zero and minimum imputation. Missing values were simulated according to the procedure described by [6] (see Methods), which introduces both missing completely at random (MCAR) and missing not at random (MNAR) values in predefined proportions. In this study, the proportions of MNAR masked values were set to 25% (left), 50% (middle), and 75% (right). For each condition, 10 different random seeds were used, and the mean MAE across these seeds is shown, with error bars representing the standard error of the MAE. The MAE was calculated exclusively for masked values based on the difference between the imputed values and the original observed values prior to masking. Distinct marker shapes indicate the MAE for only MCAR masked values, MNAR masked values, and for all masked values combined (all).

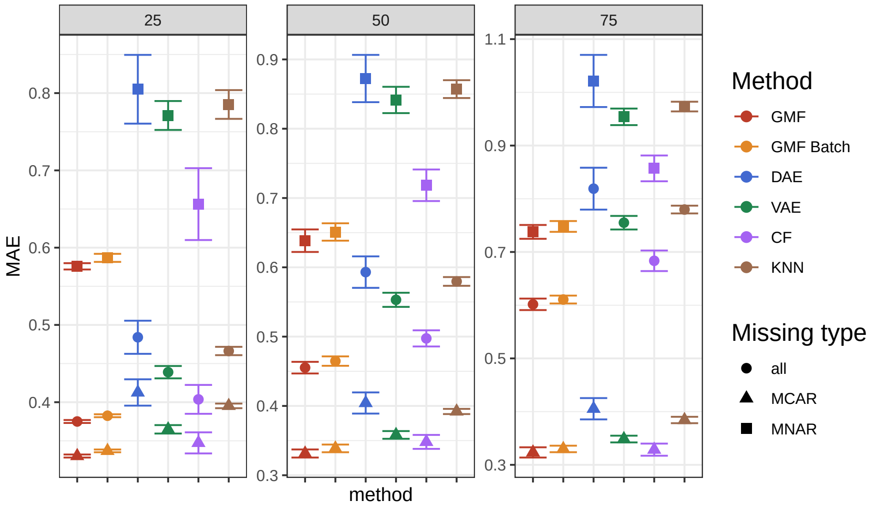


**Supp. Fig. 16** Mean average error (MAE) of the imputed values in the complete CPTAC dataset [3] evaluated for omicsGMF (GMF), omicsGMF including 2 dummy variables for the batch effects (GMF Batch), DAE, VAE, CF and KNN-imputation. NIPALS was not included due to convergence issues. Missing values were simulated according to the procedure described by [6] (see Methods), which introduces both missing completely at random (MCAR) and missing not at random (MNAR) values in predefined proportions. In this study, the proportions of MNAR masked values were set to 25% (left), 50% (middle), and 75% (right). For each condition, 10 different random seeds were used, and the mean MAE across these seeds is shown, with error bars representing the standard error of the MAE. The MAE was calculated exclusively for masked values based on the difference between the imputed values and the original observed values prior to masking. Distinct marker shapes indicate the MAE for only MCAR masked values, MNAR masked values, and for all masked values combined (all).

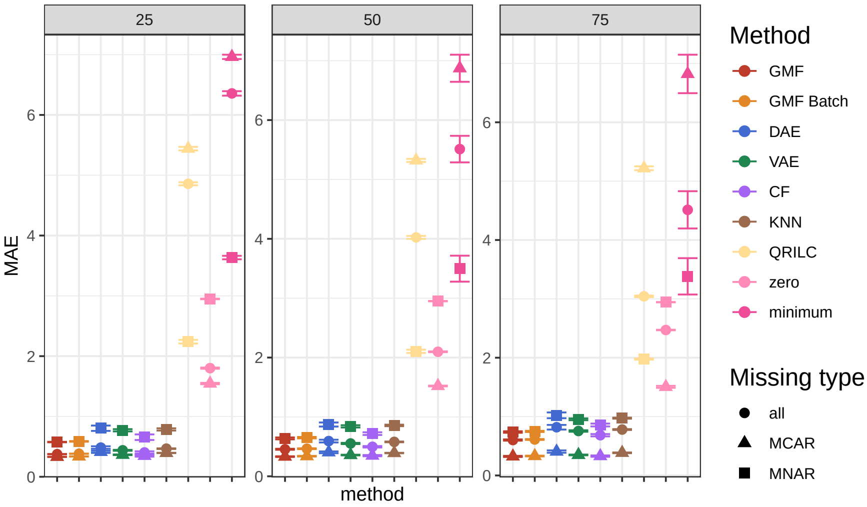


**Supp. Fig. 17** Same figure as Supp. Fig 16, but with additional methods. Mean average error (MAE) of the imputed values in the complete CPTAC dataset [3] evaluated for omicsGMF (GMF), omicsGMF including 2 dummy variables for the batch effects (GMF Batch), DAE, VAE, CF and KNN, QRILC, zero and minimum imputation. NIPALS was not included due to convergence issues. Missing values were simulated according to the procedure described by [6] (see Methods), which introduces both missing completely at random (MCAR) and missing not at random (MNAR) values in predefined proportions. In this study, the proportions of MNAR masked values were set to 25% (left), 50% (middle), and 75% (right). For each condition, 10 different random seeds were used, and the mean MAE across these seeds is shown, with error bars representing the standard error of the MAE. The MAE was calculated exclusively for masked values based on the difference between the imputed values and the original observed values prior to masking. Distinct marker shapes indicate the MAE for only MCAR masked values, MNAR masked values, and for all masked values combined (all).

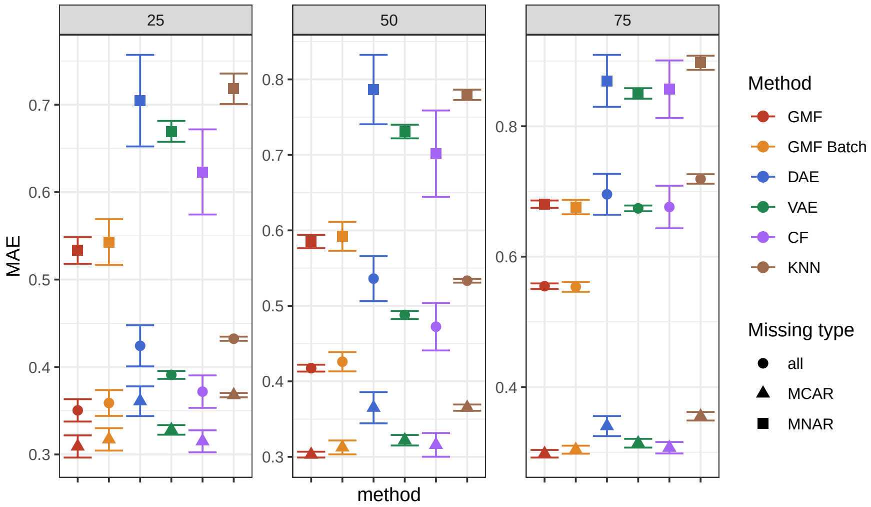


**Supp. Fig. 18** Mean average error (MAE) of the imputed values in the CPTAC dataset [3], excluding data from Lab 1 that suffered from ionization issues. The MAE is evaluated for omicsGMF (GMF), omicsGMF including a dummy variable for the batch effects (GMF Batch), DAE, VAE, CF and KNN-imputation. NIPALS was not included due to convergence issues. Missing values were simulated according to the procedure described by [6] (see Methods), which introduces both missing completely at random (MCAR) and missing not at random (MNAR) values in predefined proportions. In this study, the proportions of MNAR masked values were set to 25% (left), 50% (middle), and 75% (right). For each condition, 10 different random seeds were used, and the mean MAE across these seeds is shown, with error bars representing the standard error of the MAE. The MAE was calculated exclusively for masked values based on the difference between the imputed values and the original observed values prior to masking. Distinct marker shapes indicate the MAE for only MCAR masked values, MNAR masked values, and for all masked values combined (all).

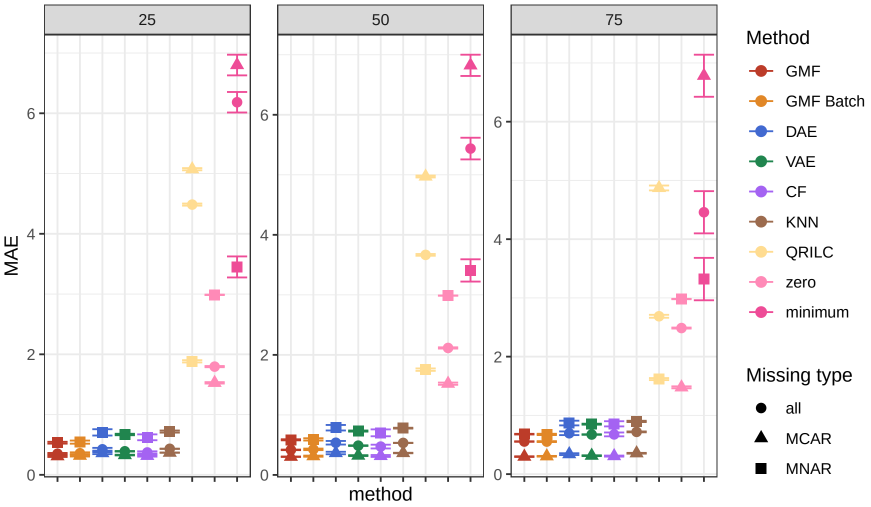


**Supp. Fig. 19** Same figure as Supp. Fig 18, but with additional methods. Mean average error (MAE) of the imputed values in the CPTAC dataset [3], excluding data from Lab 1 that suffered from ionization issues. The MAE is evaluated for omicsGMF (GMF), omicsGMF including a dummy variable for the batch effects (GMF Batch), DAE, VAE, CF and KNN, QRILC, zero and minimum imputation. NIPALS was not included due to convergence issues. Missing values were simulated according to the procedure described by [6] (see Methods), which introduces both missing completely at random (MCAR) and missing not at random (MNAR) values in predefined proportions. In this study, the proportions of MNAR masked values were set to 25% (left), 50% (middle), and 75% (right). For each condition, 10 different random seeds were used, and the mean MAE across these seeds is shown, with error bars representing the standard error of the MAE. The MAE was calculated exclusively for masked values based on the difference between the imputed values and the original observed values prior to masking. Distinct marker shapes indicate the MAE for only MCAR masked values, MNAR masked values, and for all masked values combined (all).

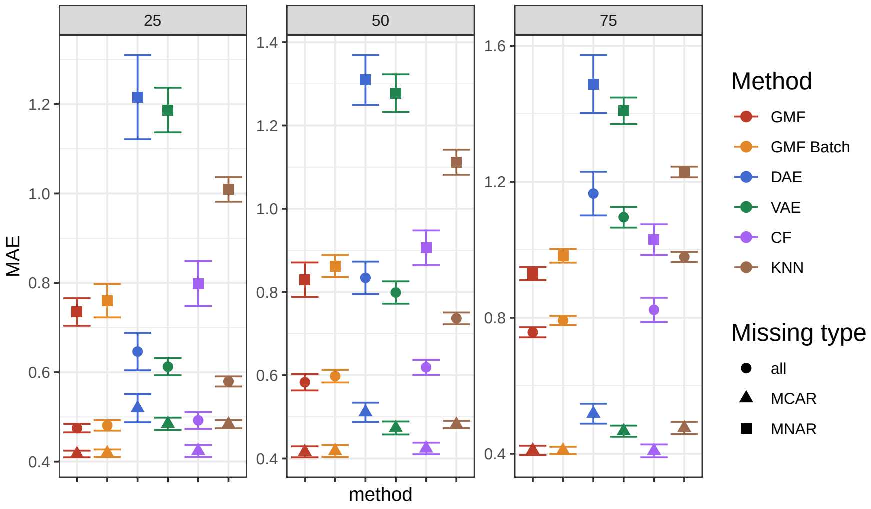


**Supp. Fig. 20** Mean average error (MAE) for the imputed values from lab 1 of the CPTAC dataset [3] when feeding all data to imputation analysis. The MAE is evaluated for omicsGMF (GMF), omicsGMF including 2 dummy variables for the batch effects (GMF Batch), DAE, VAE, CF and KNN-imputation. NIPALS was not included due to convergence issues. Missing values were simulated according to the procedure described by [6] (see Methods), which introduces both missing completely at random (MCAR) and missing not at random (MNAR) values in predefined proportions. In this study, the proportions of MNAR masked values were set to 25% (left), 50% (middle), and 75% (right). For each condition, 10 different random seeds were used, and the mean MAE across these seeds is shown, with error bars representing the standard error of the MAE. The MAE was calculated exclusively for masked values based on the difference between the imputed values and the original observed values prior to masking. Distinct marker shapes indicate the MAE for only MCAR masked values, MNAR masked values, and for all masked values combined (all).

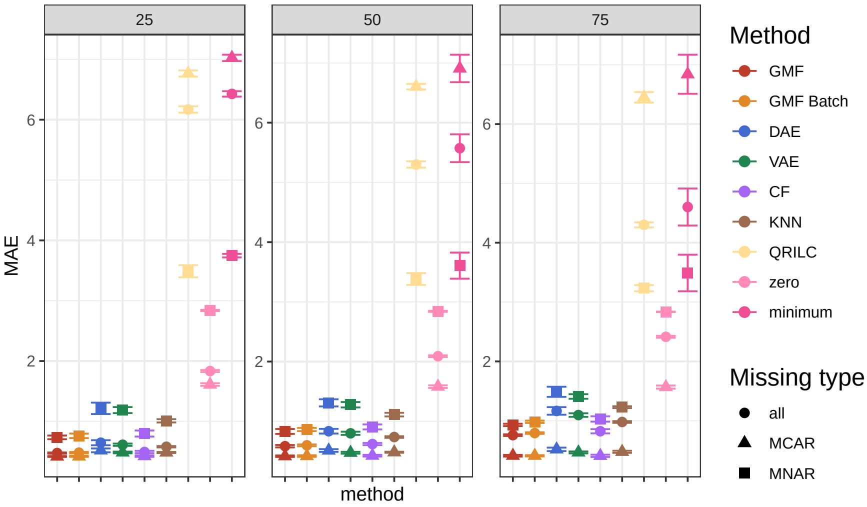


**Supp. Fig. 21** Same figure as Supp. Fig 20, but with additional methods. Mean average error (MAE) for the imputed values from lab 1 of the CPTAC dataset [3] when feeding all data to imputation analysis. The MAE is evaluated for omicsGMF (GMF), omicsGMF including 2 dummy variables for the batch effects (GMF Batch), DAE, VAE, CF and KNN, QRILC, zero and minimum imputation. NIPALS was not included due to convergence issues. Missing values were simulated according to the procedure described by [6] (see Methods), which introduces both missing completely at random (MCAR) and missing not at random (MNAR) values in predefined proportions. In this study, the proportions of MNAR masked values were set to 25% (left), 50% (middle), and 75% (right). For each condition, 10 different random seeds were used, and the mean MAE across these seeds is shown, with error bars representing the standard error of the MAE. The MAE was calculated exclusively for masked values based on the difference between the imputed values and the original observed values prior to masking. Distinct marker shapes indicate the MAE for only MCAR masked values, MNAR masked values, and for all masked values combined (all).

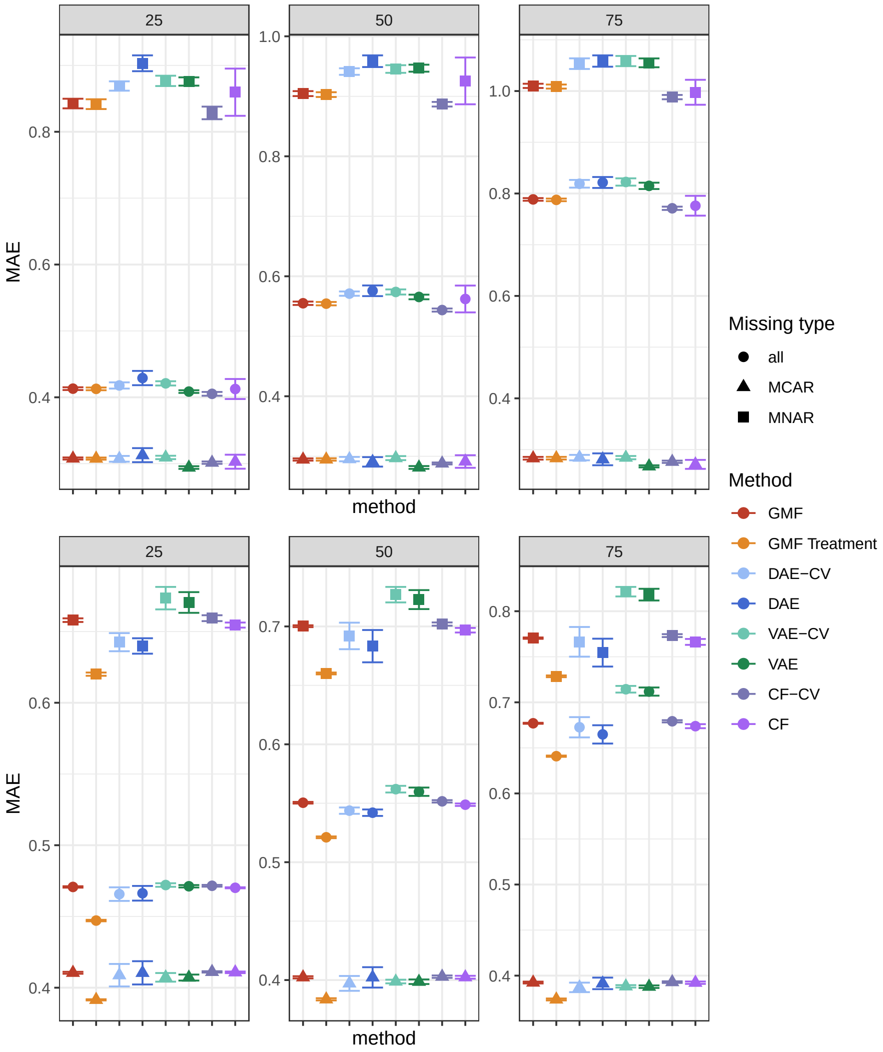


**Supp. Fig. 22** Mean average error (MAE) of the imputed values in the Petrosius dataset [1] (top) and Leduc dataset [2] (bottom) evaluated for omicsGMF (GMF), omicsGMF including a dummy variable for the treatment effect or 142 dummy variables for the batch effects (GMF Batch), DAE-CV, DAE, VAE-CV, VAE, CF-CV and CF imputation. The methods including 'CV' use the dimensionality of the latent representation suggested by omicsGMF, while the methods without 'CV' use their default values (see Methods). Missing values were simulated according to the procedure described by [6] (see Methods), which introduces both missing completely at random (MCAR) and missing not at random (MNAR) values in predefined proportions. In this study, the proportions of MNAR masked values were set to 25% (left), 50% (middle), and 75% (right). For each condition, 10 different random seeds were used, and the mean MAE across these seeds is shown, with error bars representing the standard error of the MAE. The MAE was calculated exclusively for masked values based on the difference between the imputed values and the original observed values prior to masking. Distinct marker shapes indicate the MAE for only MCAR masked values, MNAR masked values, and for all masked values combined (all).

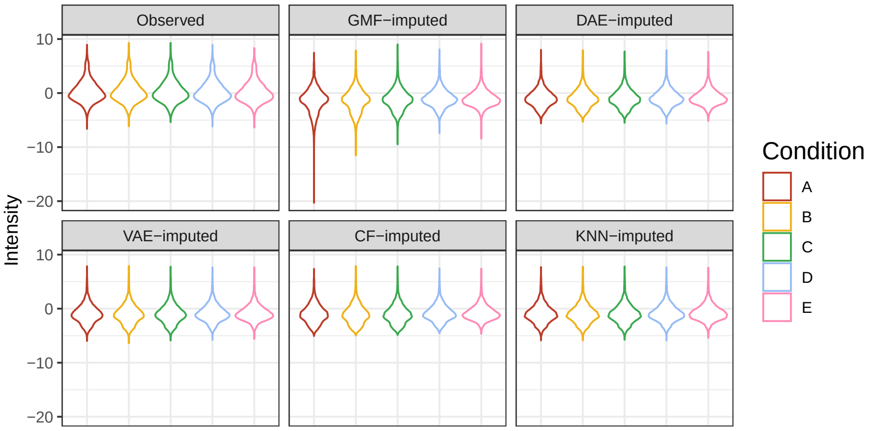


**Supp. Fig. 23** Distributions of peptide intensities for background yeast proteins of the CPTAC study, excluding Lab 1 that suffers from ionization issues, stratified according to the spike-in condition. The first panel shows the distribution of observed values, and the other panels show the distributions of imputed intensities by omicsGMF, DAE, VAE, CF and KNN imputation, respectively.

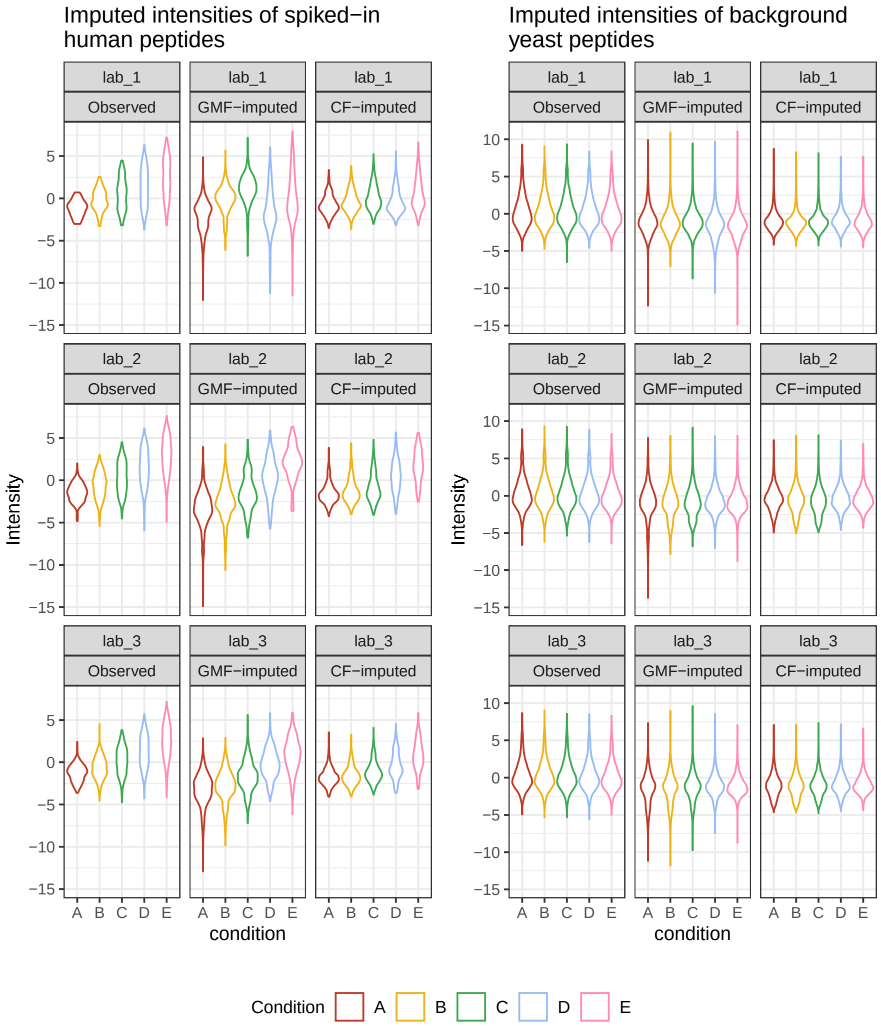


**Supp. Fig. 24** Distributions of peptide intensities for human spike-in proteins (left) and background yeast proteins (right) of the complete CPTAC study, stratified according to spike-in condition and lab. The first panel shows the distribution of observed values, and the other panels show the distributions of imputed intensities by omicsGMF and CF, respectively.

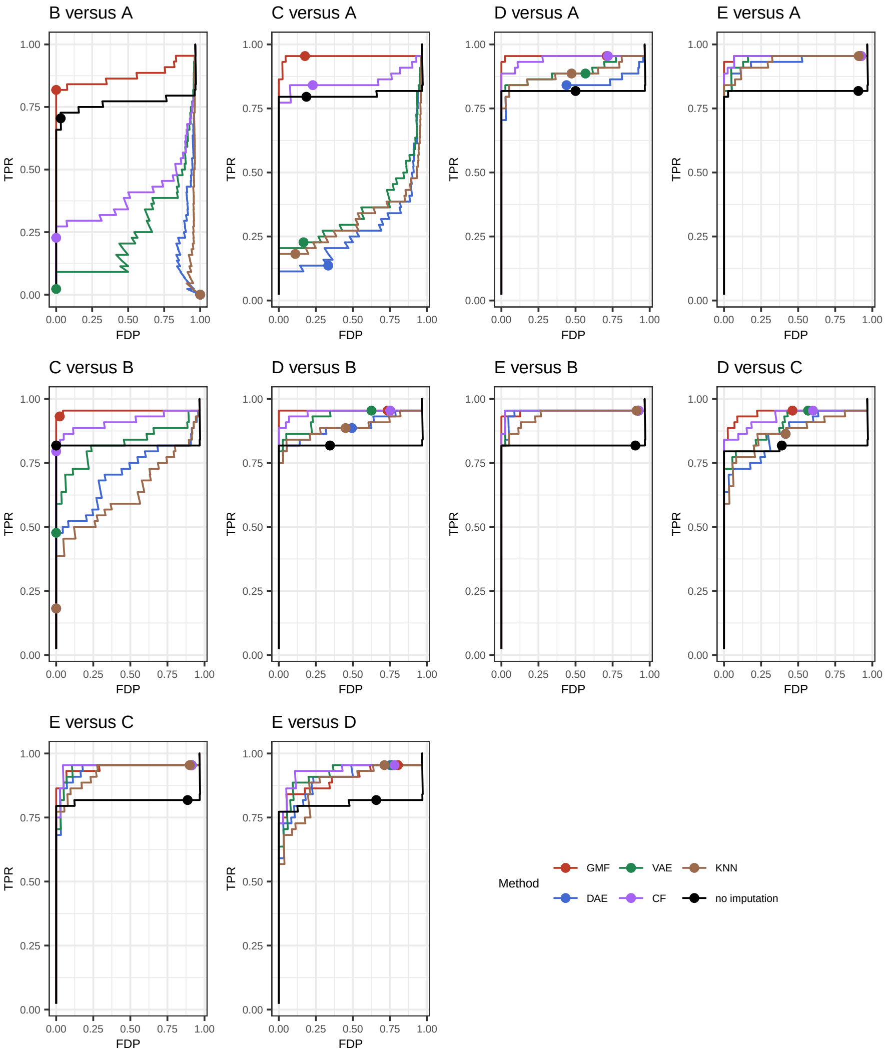


**Supp. Fig. 25** Performance evaluation of differential abundance analyses using msqrob2 [7, 8] on the CPTAC dataset [3]. Data from Lab 1 suffering from ionization issues are excluded. The human UPS proteins are differentially spiked between the conditions and the yeast background proteins serve as a true negative control. Each curve shows the true positive rate (TPR) in function of the false discovery proportion (FDP). The dots on each curve represent working points when the FDR level is set at the nominal 5% level. Results for all pairwise comparisons between the spike-in concentrations are shown.

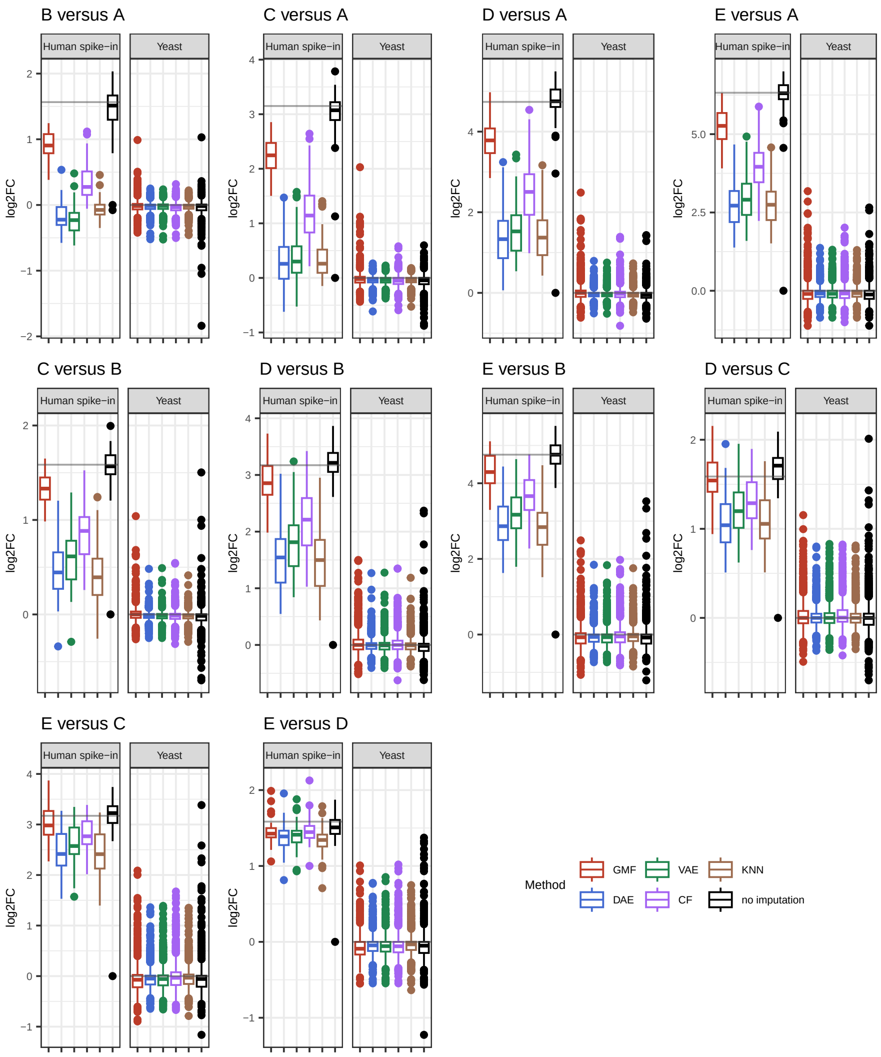


**Supp. Fig. 26** Performance evaluation of differential abundance analyses using msqrob2 [7, 8] on the CPTAC dataset [3]. Data from Lab 1 suffering from ionization issues are excluded. The human UPS proteins are differentially spiked between the conditions and the yeast background proteins serve as a true negative control. Each plot shows the estimated log2 fold changes (FC) by msqrob2 for both human spike-in proteins, and for reference yeast proteins. The grey line indicates the known log2 FC.

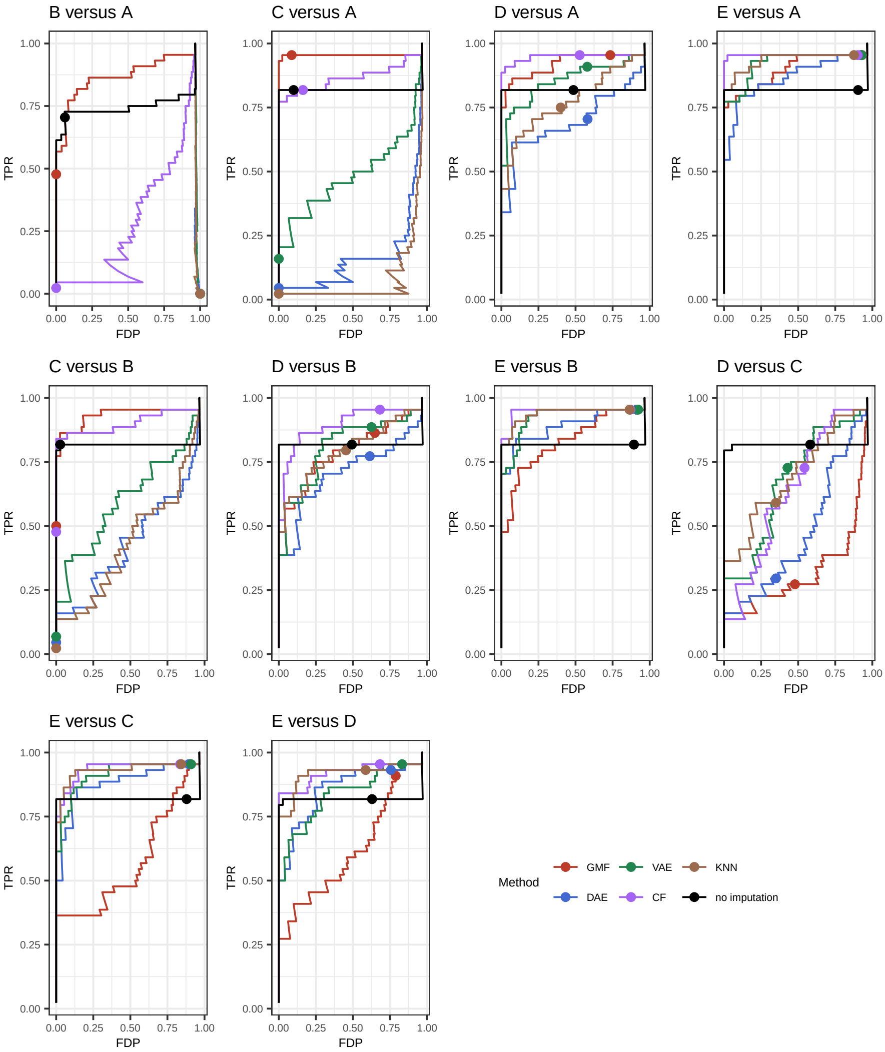


**Supp. Fig. 27** Performance evaluation of differential abundance analyses using msqrob2 [7, 8] on the complete CPTAC dataset [3]. The human UPS proteins are differentially spiked between the conditions and the yeast background proteins serve as a true negative control. Each curve shows the true positive rate (TPR) in function of the false discovery proportion (FDP). The dots on each curve represent working points when the FDR level is set at the nominal 5% level. Results for all pairwise comparisons between the spike-in concentrations are shown.

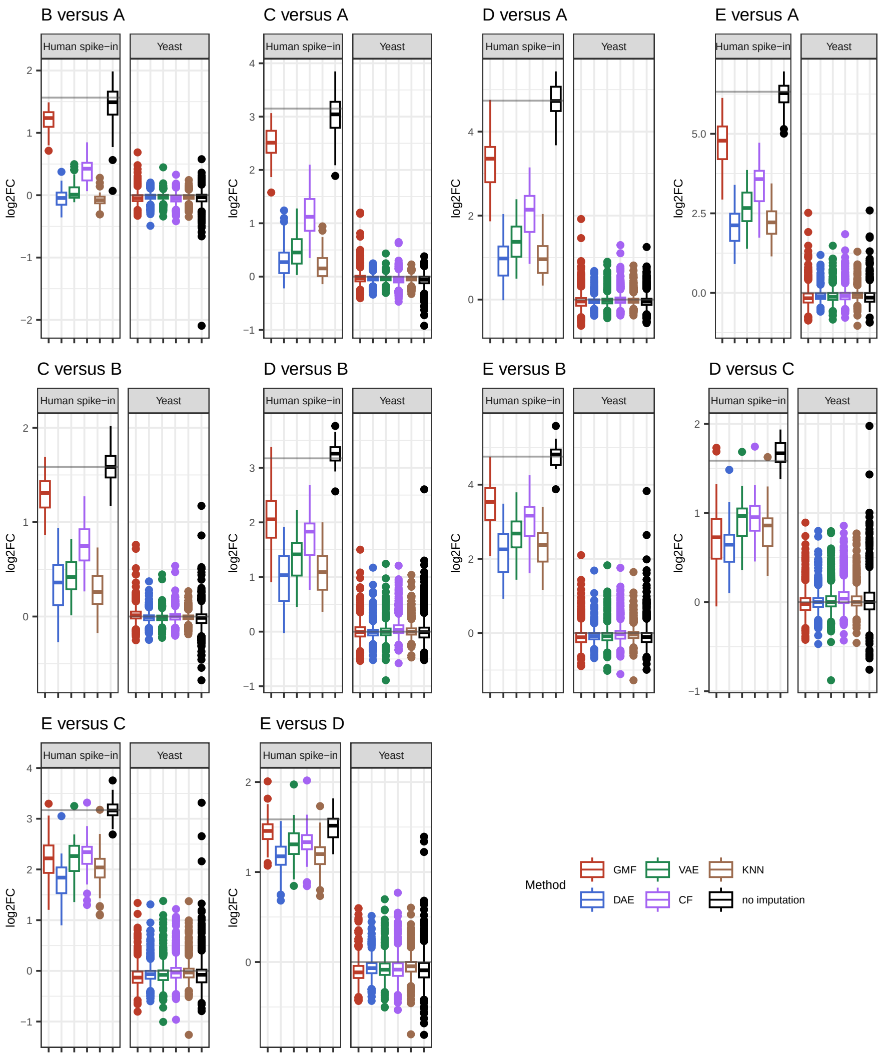


**Supp. Fig. 28** Performance evaluation of differential abundance analyses using msqrob2 [7, 8] on the complete CPTAC dataset [3]. The human UPS proteins are differentially spiked between the conditions and the yeast background proteins serve as a true negative control. Each plot shows the estimated log2 fold changes (FC) by msqrob2 for both human spike-in proteins, and for reference yeast proteins. The grey line indicates the known log2 FC.

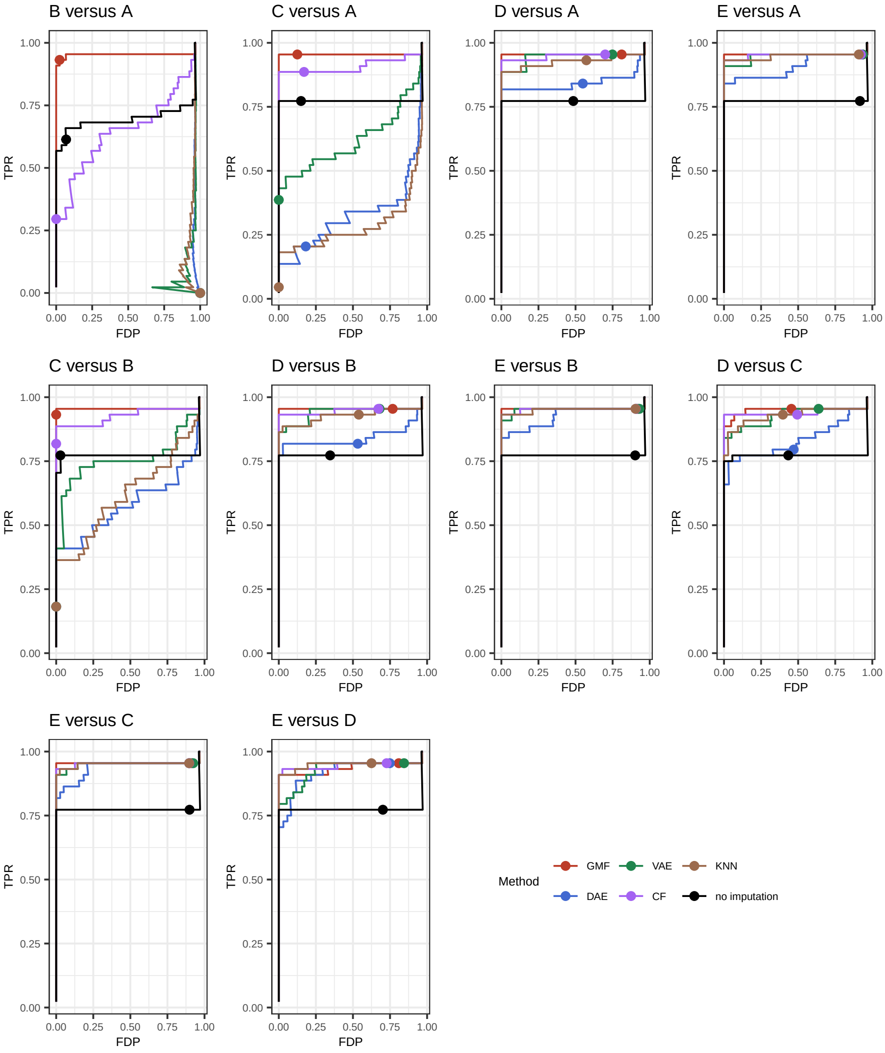


**Supp. Fig. 29** Performance evaluation of differential abundance analyses using msqrob2 [7, 8] on the complete CPTAC dataset [3], when including an additional dummy variable for conditions D-E from Lab 1. The human UPS proteins are differentially spiked between the conditions and the yeast background proteins serve as a true negative control. Each curve shows the true positive rate (TPR) in function of the false discovery proportion (FDP). The dots on each curve represent working points when the FDR level is set at the nominal 5% level. Results for all pairwise comparisons between the spike-in concentrations are shown.

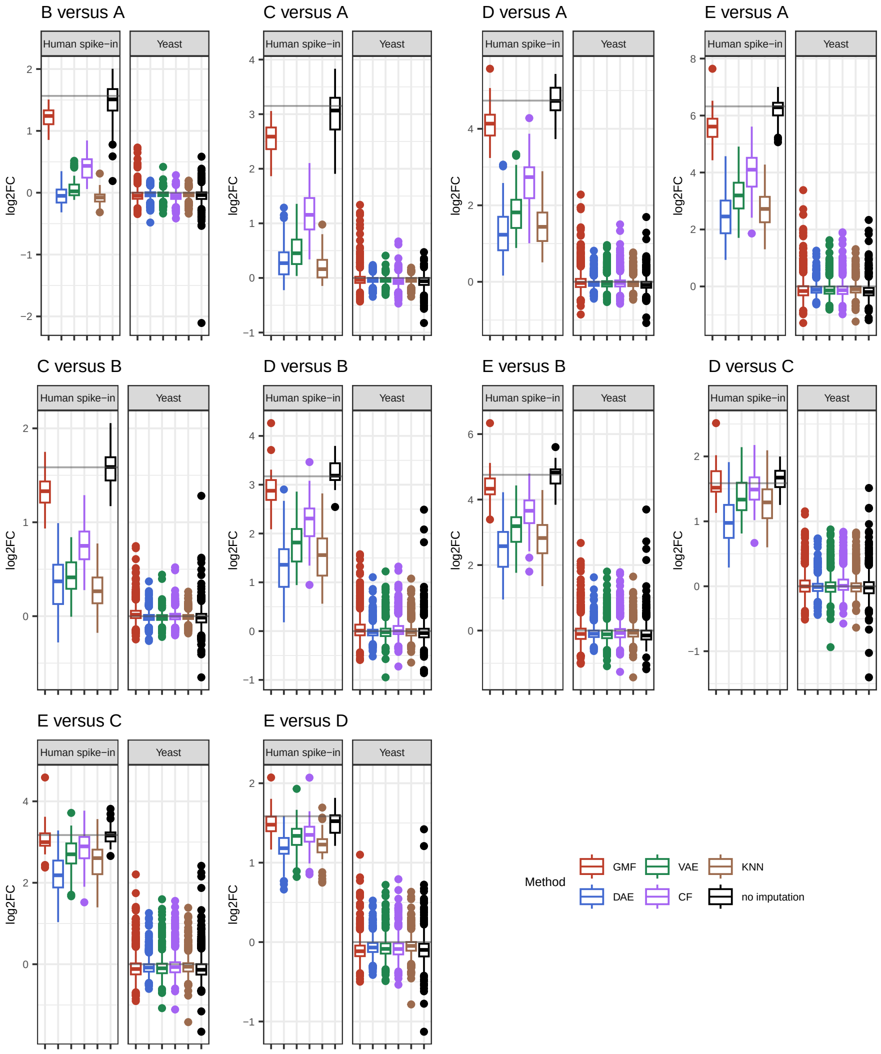


**Supp. Fig. 30** Performance evaluation of differential abundance analyses using msqrob2 [7, 8] on the complete CPTAC dataset [3], when including an additional dummy variable for conditions D-E from Lab 1. The human UPS proteins are differentially spiked between the conditions and the yeast background proteins serve as a true negative control. Each plot shows the estimated log2 fold changes (FC) by msqrob2 for both human spike-in proteins, and for reference yeast proteins. The grey line indicates the known log2 FC.

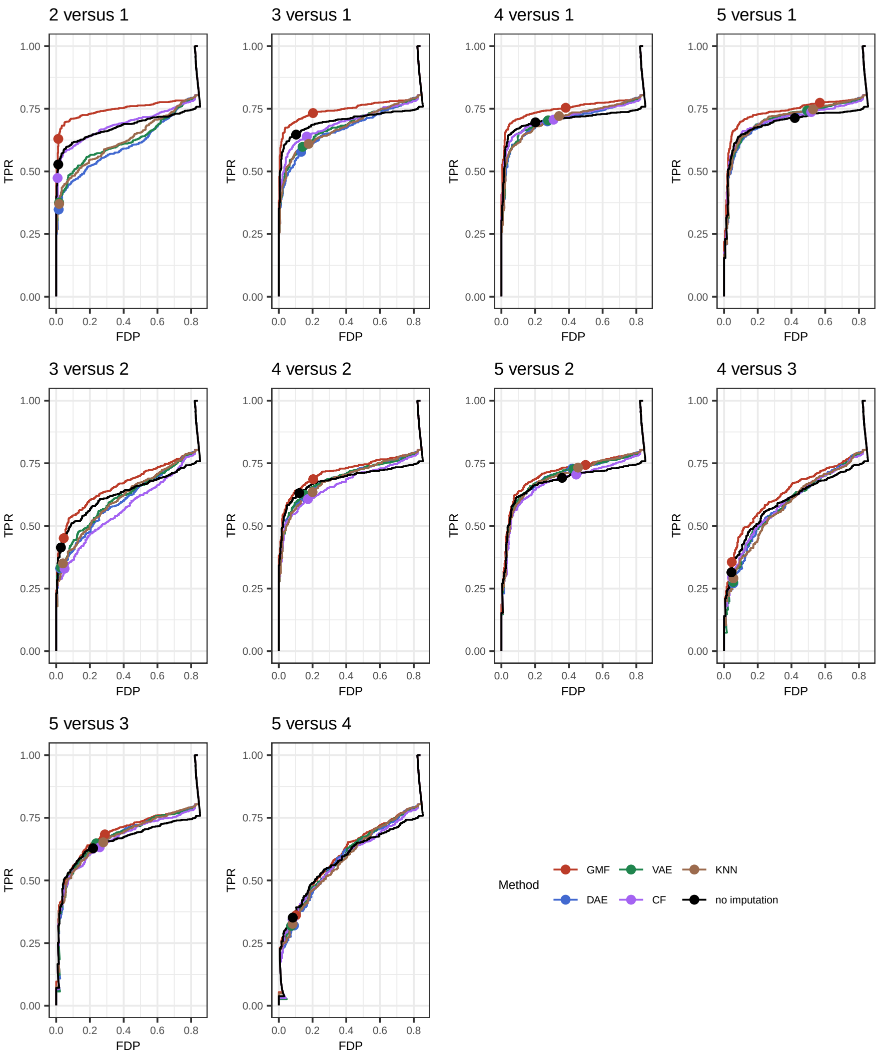


**Supp. Fig. 31** Performance evaluation of differential abundance analyses using msqrob2 [7, 8] on the Shen dataset [9]. The E. coli proteins are mixed in different concentrations within a human background. Each curve shows the true positive rate (TPR) in function of the false discovery proportion (FDP). The dots on each curve represent working points when the FDR level is set at the nominal 5% level. Results for all pairwise comparisons between the spike-in concentrations are shown.

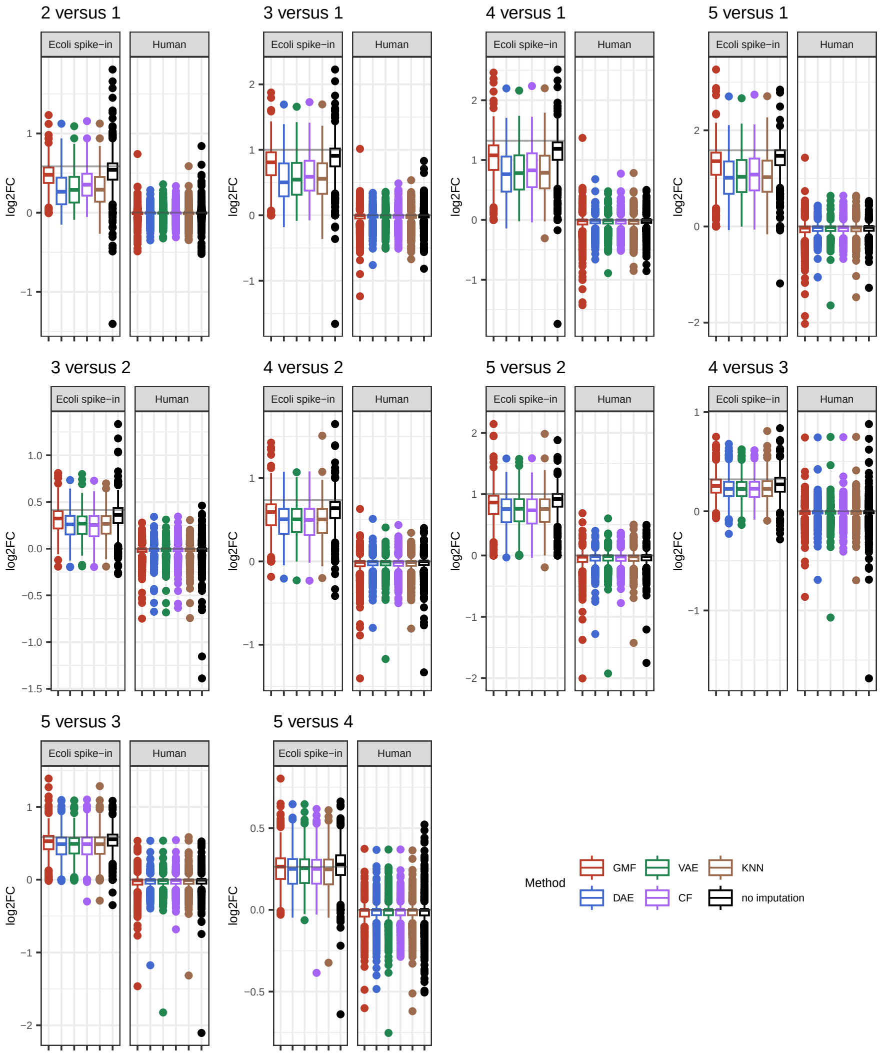


**Supp. Fig. 32** Performance evaluation of differential abundance analyses using msqrob2 [7, 8] on the Shen dataset [9]. The E. coli proteins are mixed in different concentrations within a human background. Each plot shows the estimated log2 fold changes (FC) by msqrob2 for both the spike-in E. coli proteins, and for reference human proteins. The grey line indicates the known log2 FC.

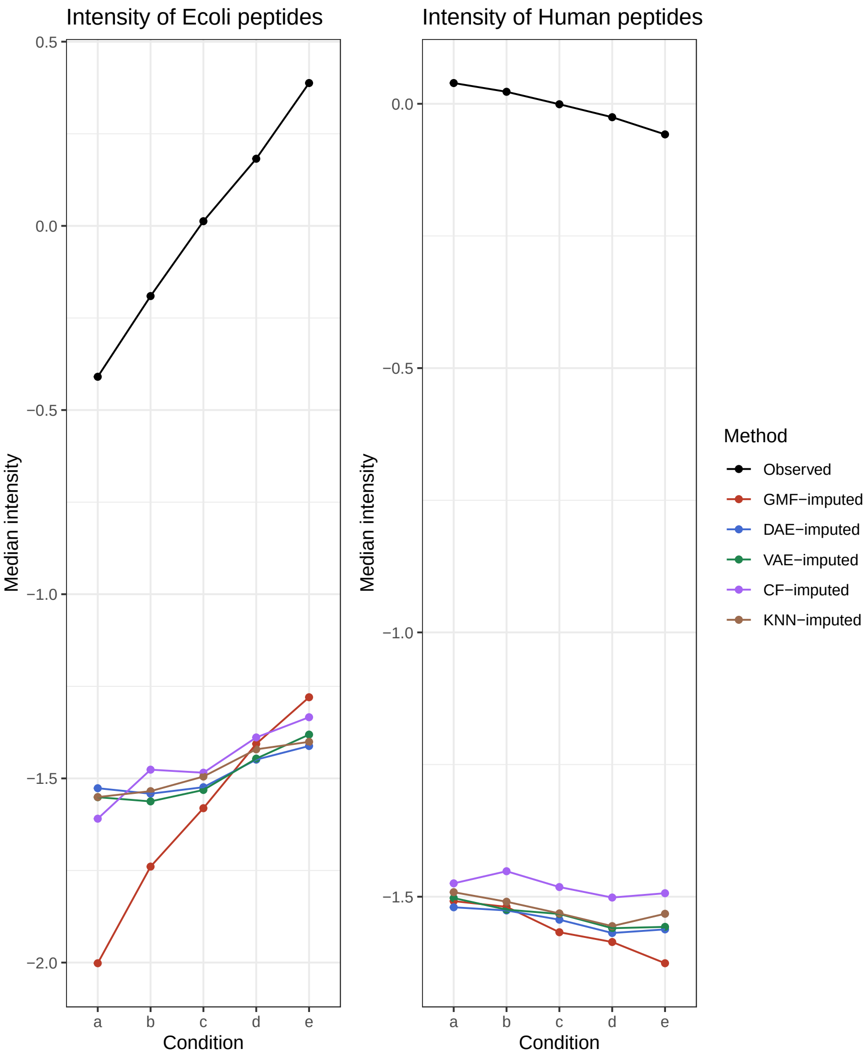


**Supp. Fig. 33** Median peptide intensities of the observed values and values imputed with omicsGMF (GMF), DAE, VAE, CF and KNN-imputation in function of the spike-in condition in the Shen study [9]. The left panel shows data from E. coli spike-in proteins and the right panel from human background proteins (right).
